# Quantifying Microbial Guilds

**DOI:** 10.1101/2023.07.23.550202

**Authors:** Juan Rivas-Santisteban, Pablo Yubero, Semidán Robaina-Estévez, José M. González, Javier Tamames, Carlos Pedrós-Alió

**Affiliations:** Microbiome Analysis Laboratory, CNB-CSIC, Spain; Logic of Genomic Systems Laboratory, CNB-CSIC, Spain; Department of Microbiology, University of La Laguna, Spain

**Keywords:** *microbial ecology*, *microbial guilds*, *phylogenetics*, *functional ecology*, *polyamine up-take*, *ammonia oxidation*

## Abstract

The ecological role of microorganisms is of utmost importance due to their multiple interactions with the environment. However, assessing the contribution of individual taxonomic groups has proven difficult despite the availability of high throughput data, hindering our understanding of such complex systems. Here, we propose a quantitative definition of guild that is readily applicable to metagenomic data. Our framework focuses on the functional character of protein sequences, as well as their diversifying nature. First, we discriminate functional sequences from the whole sequence space corresponding to a gene annotation to then quantify their contribution to the guild composition across environments. In addition, we identify and distinguish functional implementations, which are sequence spaces that have different ways of carrying out the function. We demonstrate the value of our approach with two case studies: the *ammonia oxidation* and *polyamine uptake* guilds from the Malaspina circumnavigation cruise, revealing novel ecological dynamics of the latter in marine ecosystems. Thus, the quantification of guilds helps to assess the functional role of different taxonomic groups with profound implications on the study of microbial communities.

## 1 Introduction

The large number and variety of functions carried out by microorganisms have a major impact on ecosystems. However, these functions involve a plethora of substrates and biocatalysts of varying nature like proteins [1], which are subject to evolutionary forces. Consequently, the quantitative ecological study of such functions faces several challenges.

First, proteins may diverge due to evolution, mainly through gene duplication, adaptation and drift [2– 6]. Admittedly, proteins carrying out a given function will diverge as microorganisms adapt to different conditions, while drift will cause stochastic changes between sequences despite conserving the function. These processes maintain a variety of proteins with different degrees of similarity that fulfil the same function (Fig. 1) [7, 8].

**Figure 1.**
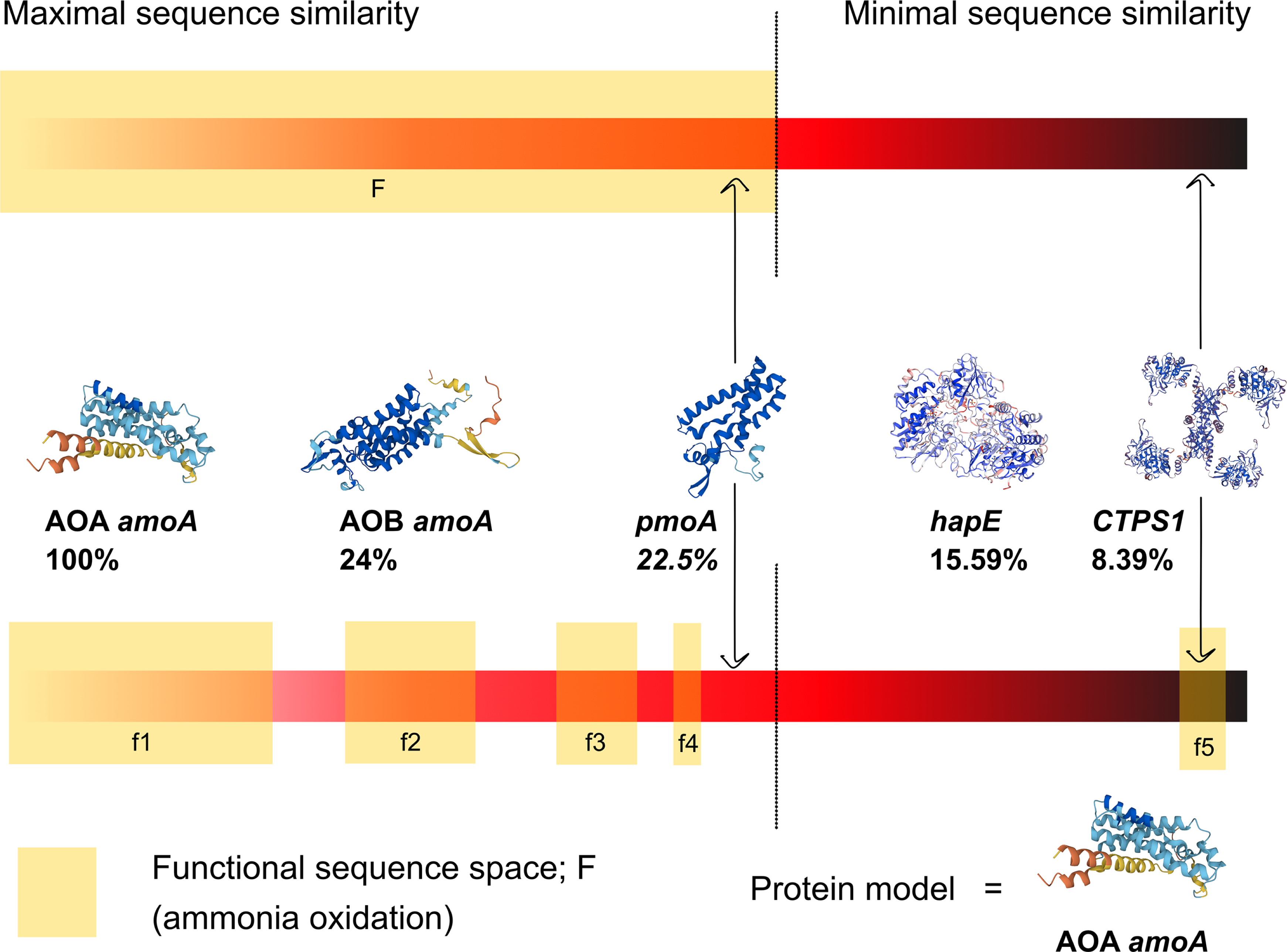
Protein sequence similarity cannot fully anticipate functional activity. Any particular function can be performed by a plethora of different protein sequences, generated by adaptive evolution and drift. Top: traditionally, automatic annotation considers any given protein sequence below a dissimilarity threshold as functional. Although the threshold can be adjusted, there are several pitfalls: (i) the threshold value is prone to errors, (ii) the functional space may display discontinuities or gaps, and (iii) proteins beyond the dissimilarity threshold may be able to perform the function. Bottom: we exemplify these scenarios with real protein sequences related to the α-subunit of the ammonia monooxygenase (*amoA*). We show: AOA (archaeal) and AOB (bacterial) *amoA* sequences well within the similarity threshold; *pmoA* which mainly oxidize n-alkanes and usually does not carry out the function but falls within the threshold; *hapE* which can oxidize a wide range of acetophenone derivatives, but not ammonia, is beyond the threshold; and CTP- synthase-1 which oxidizes L-glutamine to form a nucleic acid precursor, but it also can oxidize ammonia, thus performing the molecular function, falls well beyond any reasonable sequence similarity to AOA-*amoA*.

Second, most proteins are inherited vertically and therefore, taxonomically related organisms will often share a similar set of functions [9]. However, there are alternatives to vertical inheritance. The main one is horizontal transfer [10], which has been reported even among organisms belonging to remarkably distant lineages [11]. There are other mechanisms to acquire functions not related to inheritance, e.g. unrelated proteins converging in function [12, 13]. Therefore, neither taxonomy nor sequence similarity can fully predict the occurrence or strength of microbial functions.

There is, thus, a need for a non-taxonomic approach to analyze the ecology of microbial functions, which are mainly executed by a pool of divergent proteins. The ecological guild concept can fulfil this role. The classical definition of a guild is *“a group of species that exploit the same class of environmental resources in a similar way* (…) *without regard to taxonomic position, that overlaps significantly in their niche requirements”* [14]. This view was conceived as a way to study the ecology of macroorganisms and became popular in the 1970s. For example, all insect predators can be studied together as they are members of the *insectivorous* guild, without considering the taxonomic group they belong to [15, 16]. Thus, guilds are broadly understood as the functional groups into which communities can be subdivided, unlike the concept of population, which consists of taxonomical groups.

However, the classical definition does not suit the needs of microbial ecology. In macrofauna, guilds are usually identified by feeding behavior [17]. The range of behaviors is determined by the complex genotypic and environmental input in which an individual develops [18, 19]. Thus, the resource exploitation carried out by the *insectivorous* guild (searching, capturing, ingesting and digesting an insect) can be easily delimited and observed. In contrast, microbial feeding phenotypes are much closer to their genotypes [20]. The acquisition of a nutrient by prokaryotes depends almost exclusively on a few proteins [21, 22]. Therefore, our ability to observe microbial phenotypes is often limited to inference from sequenced metagenomes, where we rely on gene prediction and automatic function annotations, which are unreliable or simply unavailable. Furthermore, the requirement for a *similar exploitation* of resources is inherently ambiguous.

These reasons have led to a lack of consensus on how to define and quantify microbial guilds, despite a persistent use of the term [23–25]. Below are some examples.

Wu and colleagues use the term microbial guild to assign a functional value based on spatial co-occurrence among taxa [26], yet co-occurrence in space does not necessarily imply sharing the same function [27], although it might help functional prediction [28]. In another study, motility of diatoms reflected their nutritional traits and were classified accordingly [29]. However, this method is hardly extensible to all microbes. Other authors proposed that guilds should be restricted to taxa exploiting the same resource in a given space and time [30, 31], but guild membership should not be bounded spatially or temporally, since microbes from different communities may contribute to the same guild. Last, Pedrós-Alió defined microbial guilds as “*a group of microorganisms using the same energy and carbon sources and the same electron donors and acceptors*” [32]. However, this imposes a too narrow eligibility. Under this definition, methanotrophs using different electron donors while exploiting methane would not belong to the same *methane consumption* guild. It is clear, then, that different authors have understood the guild concept in different ways and this detracts from its usefulness. Could we amend such definition to better capture ecological dynamics?

Our goal is to provide a definition that fits the needs of microbial ecology, while remaining applicable to all organisms. A guild is a list of organisms that exploit or produce a key resource, and membership will be granted regardless of their lineage, their environment or how they have implemented the needed functions. Importantly, the contribution from each member to the guild varies significantly with space and time. As a result, membership is not a static quality, and the value of a member needs to be quantified in each context.

Considering this definition, we here present a method to quantify microbial guilds through the abundance, richness, and efficiency of their proteins. In addition, we improve current methods for functional delineation to strictly discriminate functional from non-functional sequences, and to categorize them ecologically. With our conceptual and methodological contributions, we avoided the problems related to similarity in niche occupancy and protein variability, which led to ambiguity and imprecision when describing microbial guilds.

## 2 Results

### 2.1 Quantification of microbial guilds from metagenomes

Our method quantifies the contribution of metagenomic sequences across three ecological aspects. Briefly, the first is the taxonomy assigned to the sequences, in our case, the protein(s); it consists of the list of guild contributors. The second distinguishes between modes in which the function is carried out, that is, how the function is implemented across the different taxa and environments. For example, *high* vs. *low affinity* transport of ammonia or *thermophilic* vs. *psychrophilic oxidation* of acetate. Often, an implementation clearly relates to taxonomy, however, in many instances a single taxon may have more than one implementation and a given implementation may be shared by several taxa (Fig. 2a). Last, we consider the environment, which allows analysis of context-dependent contributions to the guild. This approach results in a detailed picture of the guild composition (Fig. S1).

**Figure 2.**
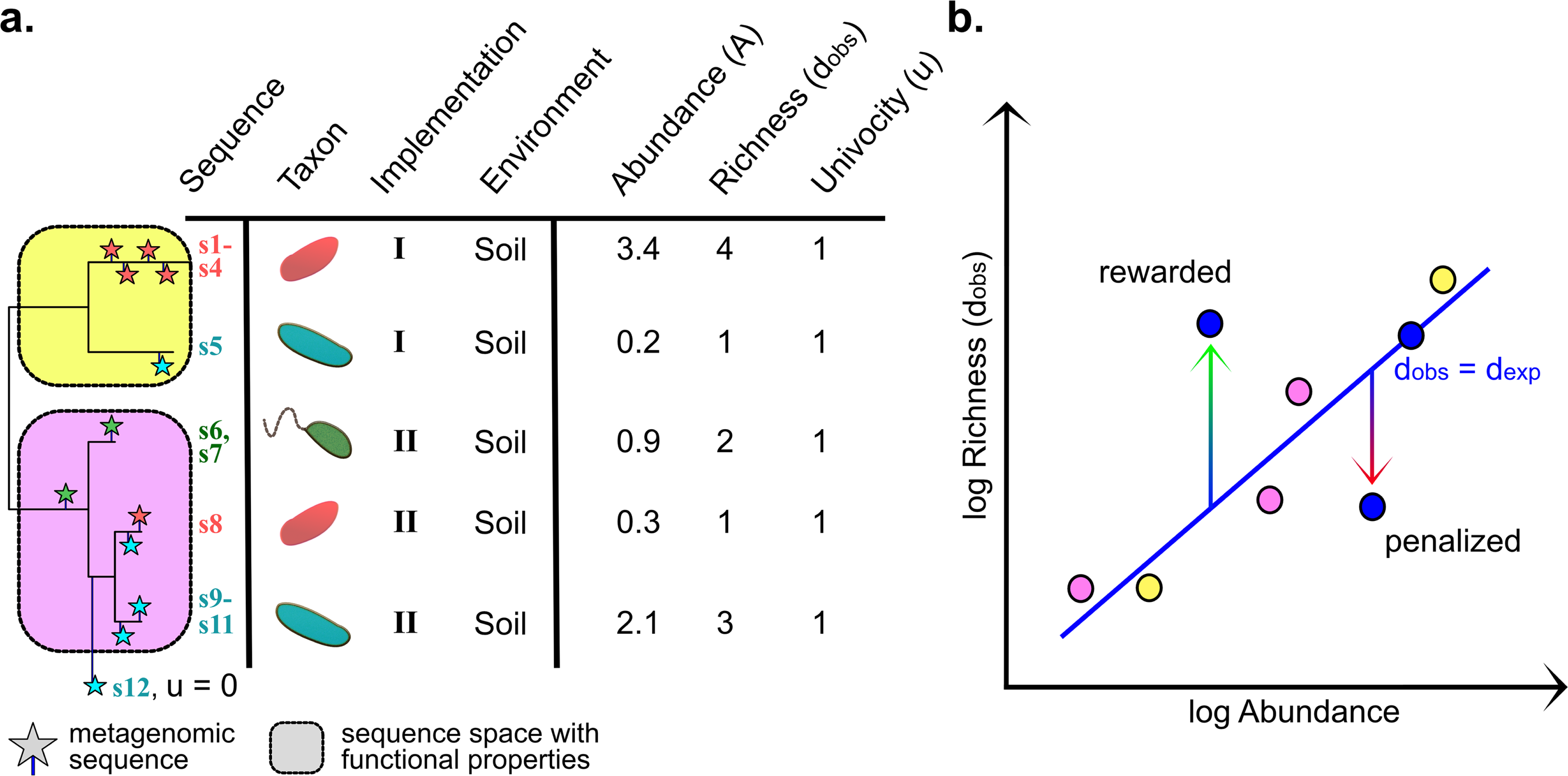
Guild quantification method. **a.** Example showing how abundance, observed diversity and univocity are calculated. A tree is constructed with reference sequences (lines, far left). Different clusters are (*u* = 1) or are not (*u* = 0) assigned to the function based on available evidence. Metagenomic sequences (stars) are placed in the reference tree. Sequence *s*_12_ does not fall within the known *u* = 1 clusters and is discarded. The remaining sequences fall in two different clusters (implementations *I* in yellow and *II* in pink). A given taxon may have different implementations and different taxa may hold the same. Next, abundance of reads for each cluster (column *A*, arbitrary abundances are shown as examples) and observed diversity (column *d_obs_*, counting the different unique sequences in each cluster) are calculated. Real metagenomic data quantification is available in Table S3. **b.** Calculation of expected diversity, *d_exp_*. In general, the more abundant a gene is the larger is the diversity of sequences. For each gene these variables show a log-log relationship. When real data from a metagenome are placed in the graph, some show either higher or lower diversity than expected from their abundance. By calculating *d_obs_/d_exp_* we reward the former and penalize the latter.

In practice, we consider that sequences associated to a taxon, an implementation and a context are equivalent, and we assess their importance considering three factors: their abundance; their richness, as the number of unique sequences; and their efficiency carrying out the function (Fig. 2a). While abundant and efficient proteins reasonably favor the function’s output, we also include richness as it typically promotes functional stability in communities and is thus of considerable importance [33], especially in microbiomes where levels of functional redundancy and niche overlap are very high [34]. In fact, it is a general postulate of ecology that the richness of the ecosystem’s constituent elements is synonymous with its complexity and maturity [35, 36]. Therefore, we quantify the contribution of a group of equivalent sequences *s* to a guild through the impact coefficient *k*:

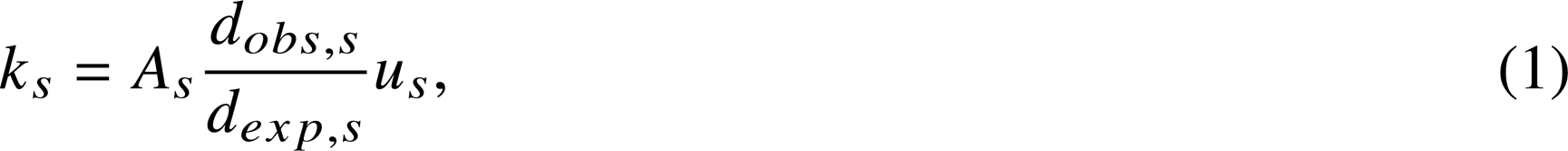

where *A* is the summed abundance of the corresponding sequences, *d_obs_* and *d_exp_* are the observed and expected richness, and *u* ∈ [0, 1] is the univocity of the function. Because we consider that the richness is key for the resilience of the function over time, we reward (or penalize) cases with higher (or lower) sequence richness than expected (Fig. 2b). The denominator *d_exp_* corrects for the fact that greater richness is invariably observed with increased abundance (Methods). Last, the univocity refers to the efficiency of the sequences. For clarity, in this study we only consider a binary classification of *u* where sequences are either functional (*u* = 1) or non-functional (*u* = 0), although this definition allows for intermediate values, for example by assessing the average efficiency to accomplish the function.

In summary, in this work we rely on normalized abundance of metagenomic reads to quantitatively characterize guilds according to taxonomy, implementations and contexts through the impact coefficients *k*, where larger values reflect the abundance in the environment of a given implementation, the occurrence of unexpected proteins richness, and the likelihood of the proteins to perform the guild-definitory function.

### 2.2 Enhancing functional profiling through the discrimination of sequence spaces in *amoA*

The previous formalism first requires a reliable functional annotation of sequences. However, automatic annotation relies on similarity with known sequences, producing many false positives because similarity alone is often not accurate enough to discriminate functionality (Fig. 1) [37]. In this section we present a method that greatly improves automatic function annotation by using reference trees as functional sequence classifiers. We illustrate the procedure using *ammonia oxidation* as an example, whose effector is typically an ammonia monooxygenase (AMO). We focus on the gene encoding the A subunit (*amoA*), which conducts the catalytic activity of the enzyme complex [38, 39] and has undergone extensive functional description.

Figure 3 shows the reference *amoA* tree to classify related sequences from our metagenomic dataset using an in-house curated oceanic database (Methods; Fig. 3a, Fig. S2 and File S1). Within the tree, we identified regions containing functional or non-functional sequences based on experimental evidence from the literature [40–42]. These regions highlight several particularities of AMO enzymes (full clustering in Fig. S3). On the one hand, *amoA* genes have evolved among taxonomic groups with different metabolic pathways. Thus, the ancestral AMO enzyme has shifted from a moderate affinity for a broad spectrum of substrates to a restricted substrate specificity, suboptimal for the substrate preference of each organism [43]. Therefore, other closely related enzymes retained some promiscuity or ambivalence; for example, particulate methane monooxygenases (pMMO) are able to oxidize ammonia even though their main substrate is methane [44, 45], or pBMO, which, apart from oxidizing butane, can hydroxylate other short alkanes (C_1_-C_9_) [46, 47]. Therefore, we can discriminate between proteins carrying out *ammonia oxidation* with *u* = 1 and others that have higher affinities for methane or other simple aliphatic alkanes [48], with *u* ∼ 0 for *ammonia oxidation* (Fig. 1). On the other hand, there are two different implementations among functional sequences: the archaeal and bacterial AMO, AOA and AOB respectively. Each occupies different niches [49] and act differently on the key substrate: AOA-*amoA* proteins display a ≈ 200-fold higher affinity for ammonia, dominating in oligotrophic environments [50].

**Figure 3.**
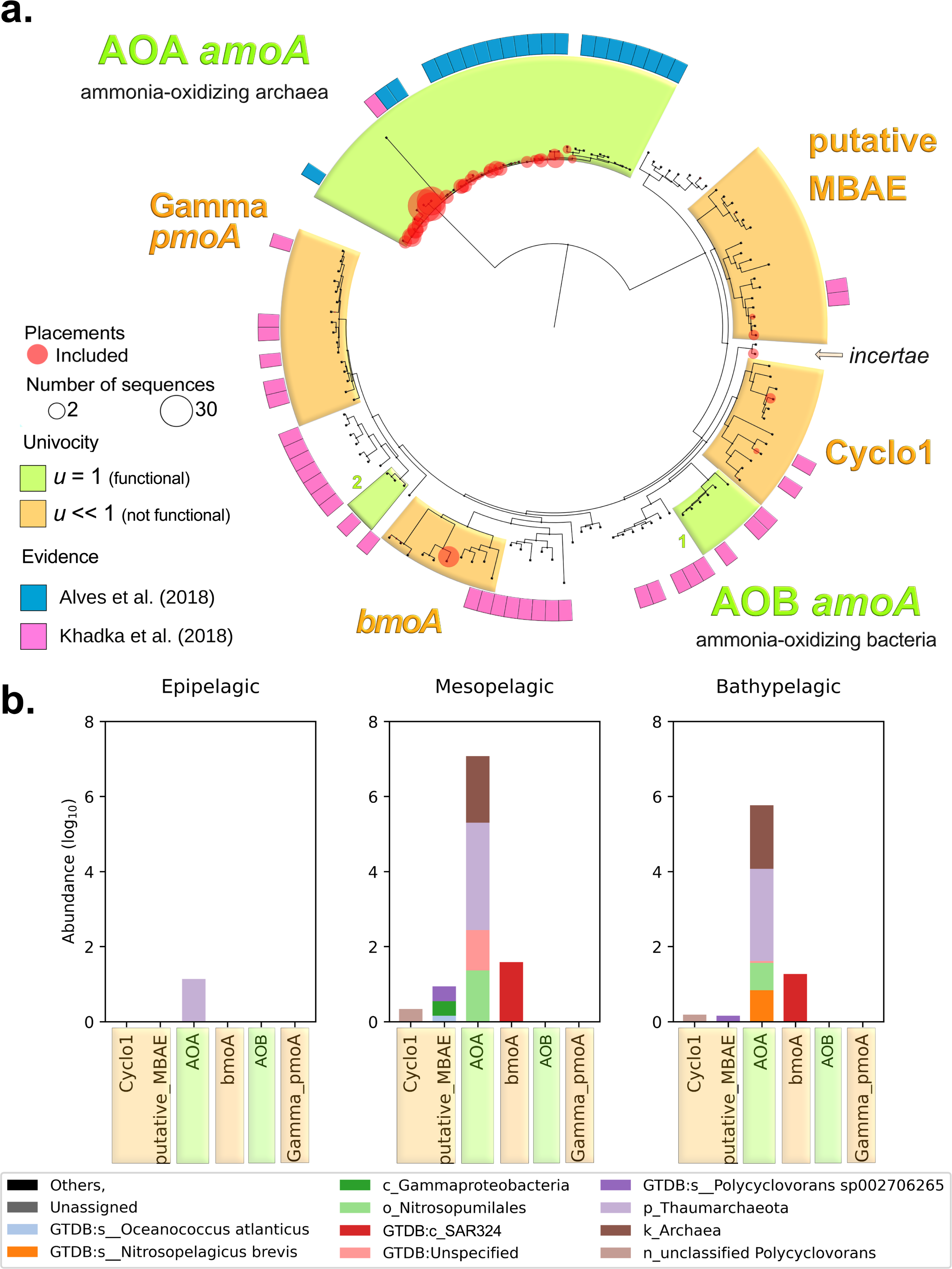
Distinguishing functional from non-functional sequences improves the assessment of the *ammonia oxidation* guild. **a.** Reconstructed phylogeny of *amoA*, used as a reference tree to classify ammonia-oxidizing capable sequences. The tree contains 135 sequences with strong functional evidence based on either biochemical or physiological features, or inferred by homology to quality sequences (see Methods). For clarity, we only highlight clusters of sequences where metagenomic Malaspina samples have been successfully placed (full clustering in Fig. S3). Among those, we distinguish sequence clusters with proven ammonium oxidation function (shaded greens) from sequences with a broader substrate spectrum and sequences with evidence of being non-functional for ammonia oxidation (shaded orange). Note that AOA-*amoA* form a single cluster, but AOB-*amoA* comprises two dissimilar clusters. Evidence of function was gathered from various sources, albeit the main ones are marked in pink and blue in the outer circle. The size of red circles is proportional to the number of reads. **b.** Log representation of abundance values (TPM) of the *amoA* classified queries found in Malaspina metagenomes (red circles from a.). We excluded 31% of unique sequences (1.01% of total TPM) corresponding to the non-univocal tree clusters, keeping a conservative criterion.

Then, we retrieved *amoA* sequences from Malaspina metagenomes [51] and placed them onto the tree (Fig. 3a). First, the placements were robust as we did not find any spurious sequence. However, we found that from a total of 129 unique sequences, 40 were discarded by the tree placement due to paralogy (31.0%), and only 82 were *bona fide* for ammonia oxidation (63.5%). Note that, despite the similarity between sequences in the AOB-*amoA* and *u* = 0 clusters, we were able to distinguish functional from non-functional proteins. In Figure 3b we show the results of the classification of the Malaspina queries per implementation, taxon and water column depth in terms of their abundance. We are able to focus our analysis on the AOA and AOB implementations (in green) which are the reliable functional sequences in contrast to existing approaches that are unable to filter out false positives, and to distinguish implementations.

Overall, this shows that several *amoA*-like sequences were non-functional, highlighting the need to recognize false positives (*u* = 0) to accurately quantify the ammonia oxidation function in metagenomes. In addition, identifying different regions in the tree allowed us to identify different *amoA* implementations, providing a more detailed description of the *ammonia oxidizers* in the Malaspina metagenomes.

### 2.3 Discriminating functional sequence spaces of *potF* based on environmental preferences

Unlike *amoA*, unfortunately, most genes have limited or ambiguous experimental evidence. This is the case of *potF*, which is a subunit of an ATP-binding cassette (ABC transporter) that binds and imports putrescine-like polyamines. Here, we were unable to delimit sequence spaces by substrate preference due to the scarcity of experimental evidence (Fig. S4), but we were able to characterize clusters using a different criterion.

As in the previous case, we built a reference tree (Fig. 4a), but using the Hidden Markov Model (HMM) corresponding to *polyamine binding*, KEGG K11073 [52] (File S2) to identify functional sequences with the same oceanic database. Thus, the reconstructed reference tree showed a collection of HMM-retrieved *potF*-like sequences grouped by their similarity where each sequence is represented by a leaf of the tree. The ability of a protein to perform a function is influenced by its surrounding environment, thus requiring specific conditions to perform it effectively. In particular, periplasmic transporters are more exposed to changing environments than cytoplasmic proteins. Therefore, we wished to explore whether the divergence between tree branches depended on environmental conditions.

**Figure 4.**
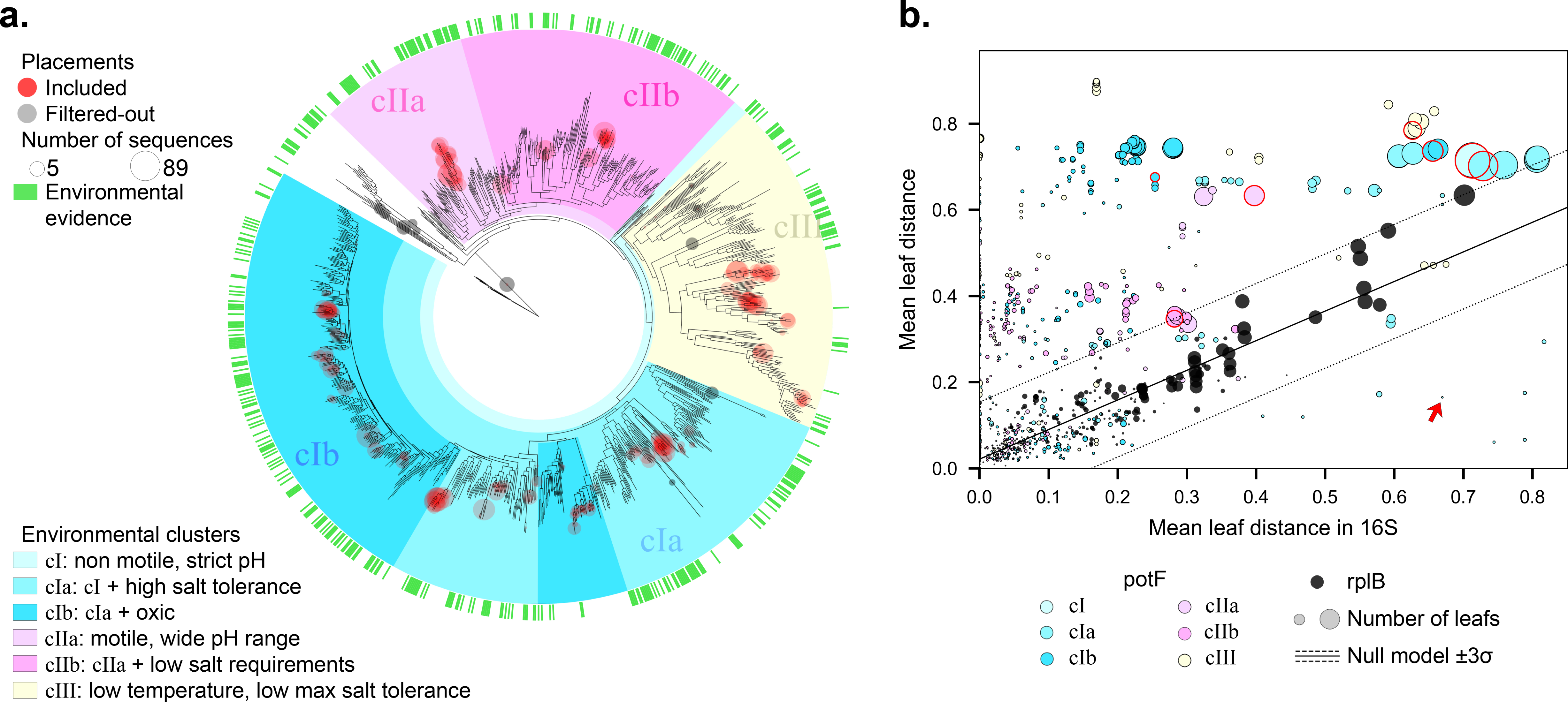
Polyamine uptake implementations revealed by environmental cues are non-synonymous with phylogeny. **a.** A phylogenetic reconstruction of *potF*, a polyamine-binding subunit of an ABC transporter. Due to the very scarce experimental evidence available, we here determine the different implementations as sequence spaces adapted to specific environmental conditions (Methods, Tables S1 & S2). We gathered environmental data for 478 tree locations representing 321 organisms (green tags). We found six nodes enriched by environmental preferences (> 10^4^ randomizations, one-tailed p-values < 0.003) which included most sequences. Thus, the tree acts as a metagenomic query classifier. **b**. To find whether the previous clusters grouped taxonomically related species, we tested the divergence between last common ancestors (LCAs) in the trees of *potF* and a pair of phylogenetic markers: ribosomal 16S RNA and *rplB* genes. The average phylogenetic distances between common groups of organisms (nodes in the tree) in the 16S RNA (x-axis) are compared to those in the *rplB* gene (black dots) or to those in the *potF* gene (colored dots). The size of the dot represents the number of children leaves compared. The two phylogenetic markers follow a straight line as expected. The *potF* sequences, on the other hand, significantly depart from this linearity. The null model is equivalent to the neutral variation expected from phylogeny and was fitted with a bivariate regression (mean and ±3σ, black solid and dotted lines). Red edges to the circles highlight the nodes that define the environmental clusters in panel a, which are beyond the taxonomy-related null model. The red arrow shows an example that corresponds with *Oceanobacter kriegii* and *Thalassobius gelatinovorus*, where their *potF* sequences are more similar than what is expected by their taxonomic markers.

To that aim, we retrieved from the literature the environmental preferences of the bacteria present in the reference tree. We obtained fifteen environmental variables for 321 organisms in pure culture, corresponding to 478 of 1158 tree leafs (Tables S1 and S2, and Fig. S5). We then searched for sequence spaces that shared similar environmental properties (randomized tests, one-tailed p-values < 0.003). First, we observed significant nodes that grouped only a handful of sequences, highlighting properties that the taxonomy accurately predicts, e.g., hydrocarbon degradation [53]. Second, we found significant nodes that grouped all sequences in just a few clusters under a combination of preferences to temperature, salinity and pH, suggesting groups undergoing either common environmental adaptation, or horizontal gene transfer (shaded colors in Fig. 4a).

We wondered whether these environmental features were deducible from taxonomy or not. To answer, we compared the tree structures between a phylomarker and *potF*-like sequences for the same list of organisms (Methods). We found that the significant internal nodes grouped sequences that diverged differently than expected from the phylomarker (Fig. 4b). This divergence can be exemplified by two different limit cases. On the one hand, the same organism may have more than one sequence, which may be either in distant or nearby positions in the tree, e.g., *Pseudomonas alcaligenes* appeared in six distant positions among three different clusters (Fig. S6). On the other hand, sequences from distant taxa may unexpectedly converge in similarity. This is the case of *Oceanobacter kriegii* and *Thalassobius gelatinovorus*, an alpha- and gamma-proteobacteria, respectively, whose normalized phylogenetic distance on a 16S tree is large, while being remarkably close in the *potF*-like tree (0.67 *vs*. 0.16, red arrow in Fig. 4b). Thus, phylogenetic signal poorly predicted divergence in *potF*-like sequences. Therefore, we used the significant internal nodes to define different implementations and classify sequences, underlining broader trends and properties across the reference tree of *potF*.

Finally, we placed the *potF*-like short queries from Malaspina metagenomes onto the annotated tree, discarding 71 (4.13%) poor quality unique sequences (Methods; grey circles in Fig. 4a). The rest of the placements had a mean weighted likelihood robustness of 0.89 and populated the entire tree, suggesting that most known marine polyamine-like binding proteins were well represented in our database. In this way, we decoupled functional behaviors from taxonomic positions of *potF*-like proteins, and discretized the sequence space using environmental evidence

### 2.4 Decoding ecological dynamics in the *polyamine uptake* guild in marine contexts

We finally quantified the *polyamine uptake* guild in the Malaspina circumnavigation samples, showcasing the potential of our approach. This specific guild is reported to be ubiquitous in the oceans [54], so it was ideal to assess the guild composition in different marine environments. Using the classified high quality sequences from the previous section (red circles in Fig. 4a) we calculated the impact coefficients *k* for each taxon, implementation and environment and analyzed the guild composition across three different depth layers: epipelagic (0 – 200 m), mesopelagic (200 – 1000 m) and bathypelagic (1000 – 4000 m).

Figure 5 compares the results between state-of-the-art methodologies (a), our approach considering only abundance (b), and our approach based on impact coefficients (c). First, note that current methodologies collapse the whole *potF* sequence space into a single bar, while we are able to differentiate between implementations. Second, the impact coefficients (which include the richness ratio, Eq. 1) reveal taxa that have a huge effector richness and that might be fundamental for functional stability, e.g., *Sulfitobacter sp001629325* and *Pseudooceanicola atlanticus*. Therefore, our approach provides a more comprehensive understanding of the guild structure and its differences across contexts.

**Figure 5.**
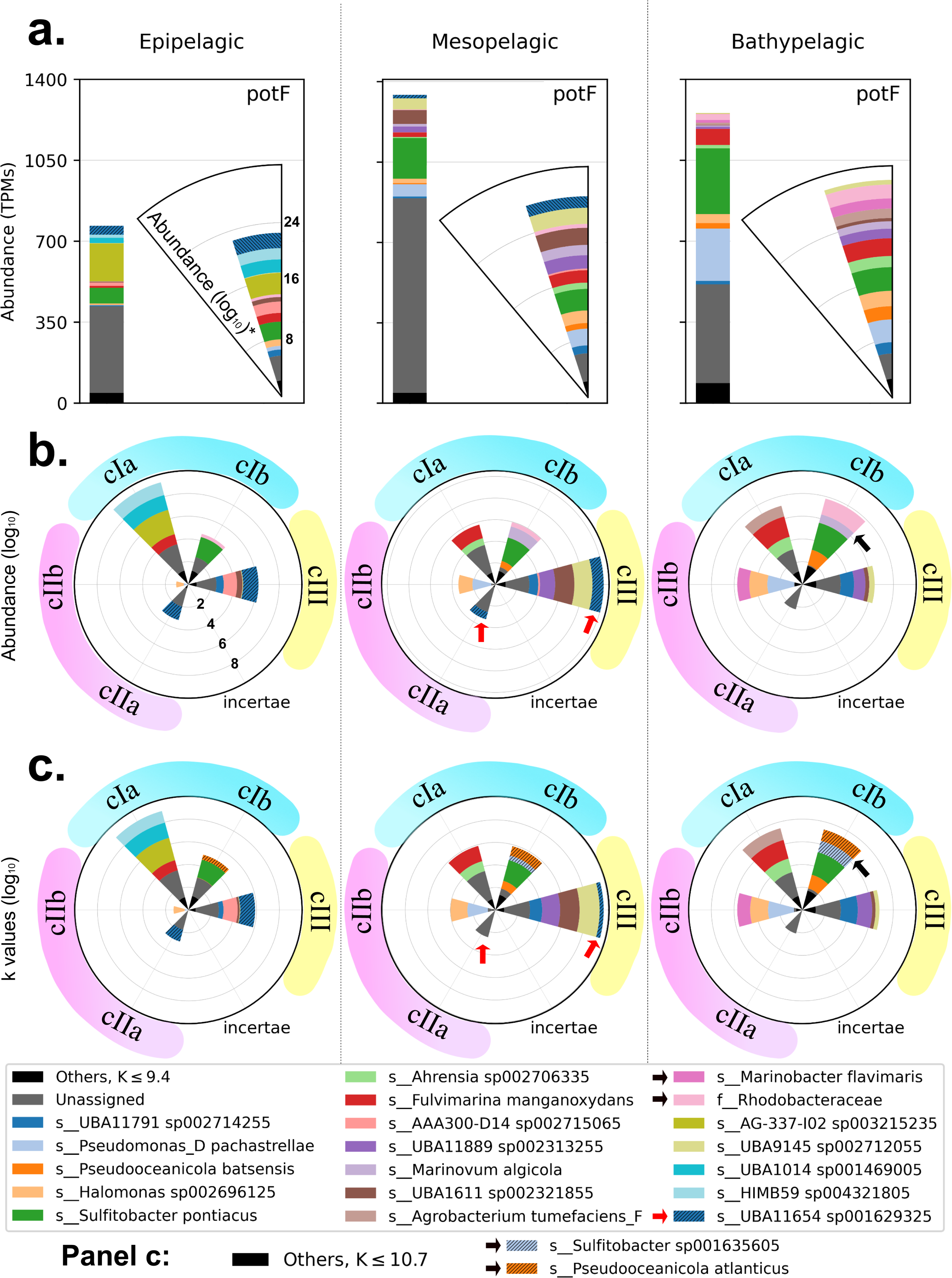
Our guild approach identifies important ecological patterns in the *polyamine uptake* guild in contrast to the traditional approach. We show (a) the output of a standard automatic metagenomic analysis. The results have been displayed as histograms and as radial plots with stacked log values to improve the visualization of taxa spanning different orders of magnitude; (b) our guild approach considering only abundance; and (c) the impact coefficients, which also ponder sequence richness; across three environments (columns). What in the automatic annotation was a single bar has been split into several sectors, each of them corresponding to one of the implementations as identified in Fig. 4a. We only show taxa with the largest contributions, the rest are grouped under *Others* tag (black). The contribution to the function fluctuates in both taxonomic identity and implementation preference with nontrivial relationships with depth. For example, UBA11654 sp001629325 (striped blue, red arrows) contributes in the epipelagic with two different implementations, *cIII* and *cIIa*, while it only contributes through *cIII* in the mesopelagic, and disappears in the bathypelagic. Moreover, *Sulfitobacter* sp. and *P. atlanticus* (black arrows) were only visible considering expected richness. Note how easy it is to observe distinct functional trends for each taxon, in contrast with traditional approaches (a). The *incertae* implementation is representing the absence of *k* values in the undefined sequence spaces of the reference tree.

Overall, our results showed that the *polyamine uptake* guild is important throughout the entire water column, but with a different composition of taxa and implementations. The main implementations of *polyamine uptake* found were those significant for activity under saline conditions (*cIa*, *cIb*, and *cIII*), which is coherent with the fact that the samples were all marine. Also, we observed considerable functional redundancy as there were different implementations in every sample and several taxa within each implementation. Specifically, the guild structure changed significantly between the epipelagic and mesopelagic, both in taxonomic composition and in the estimated strength of each of the implementations. Between the mesopelagic and bathypelagic, the pattern was remarkably taxon-preserved but the net contribution of each implementation to the overall function changed slightly, with more top contributors in the latter. However, the *polyamine uptake* function persisted throughout the water column despite these changes in taxonomic composition, showing a species turnover with depth.

In addition, we easily compared the guild across environments by aggregating implementations and/or taxa (in *s* of Eq. 1). Thus, we computed the fold-changes of the impact coefficients *k_meso_/k_epi_* = 1.82, and *k_bathy_/k_meso_* = 0.95. Even though several implementations were more important and taxonomically diverse in the bathypelagic, the main guild activity occurred in the mesopelagic. Similarly, we could also compare the guild’s composition across environments and implementations by aggregating all sequences corresponding to different taxa (in *s* of Eq. 1). The most remarkable change corresponded to implementation *cIIb* between the epipelagic and mesopelagic *k_cIIb,meso_/k_cIIb,epi_* = 11.3 (Fig. 6), suggesting an implementation-dependent bloom in the mesopelagic. This is an interesting finding, since the implementation *cIIb* is significant for organisms with a wide pH range (Fig. 4), and the greatest variability of oceanic pH occurs precisely in the mesopelagic (Fig. S7). This confirms the ability of our approach to reveal environmental-dependencies of biological functions in general.

**Figure 6.**
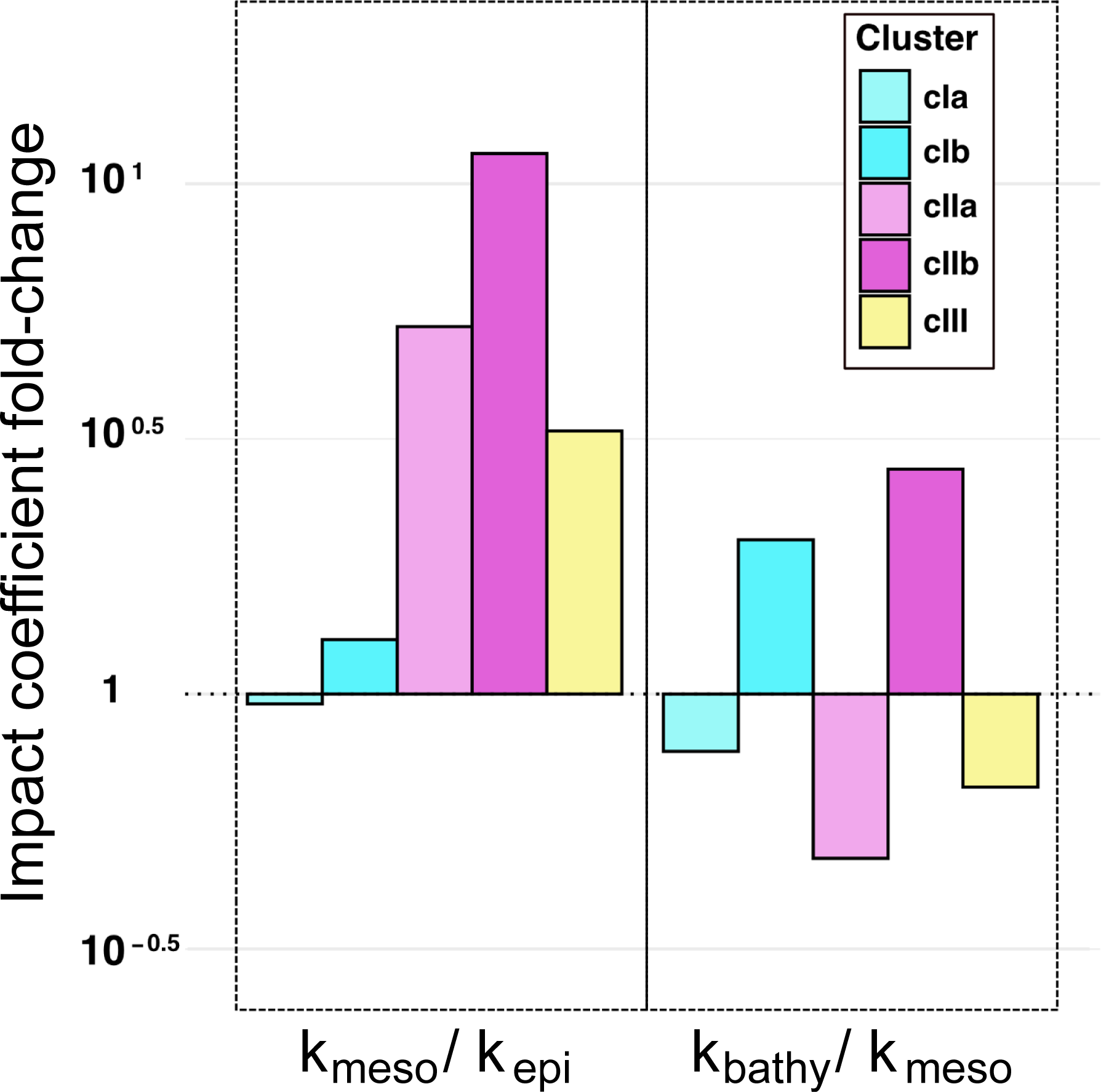
Changes with depth in the importance of the *polyamine uptake* guild. An advantage of using the impact coefficients to determine the structure of a guild is that we can visualize the functional contribution in a variety of ways. For example, here we look at the fold changes in the contribution between different ocean layers for different implementations. First, the *k* values of each implementation are added for each layer (*k_epi_, k_meso_, k_bathy_*). Then the values for two different layers are divided (*k_meso_/k_epi_*, *k_bathy_/k_meso_*). It is easily observed which implementations depend the most on depth, which in this case are *cIIa* and *cIIb*, sequence spaces putatively adapted to a wide pH range. Interestingly, the sharp changes in these two implementations correspond with the area of the water column with the largest shift in pH toward acidity in the oxycline (Fig. S7).

## 3 Discussion

### 3.1 Diversification of functional protein sequences

Most functions are performed by evolving proteins. In addition, each function is often found in many different environments. Thus, diversification of the function-capable sequences is not only expected but frequently observed [55–57]. In order to quantify microbial guilds, the issue of how proteins diversify while maintaining function must be considered.

First, diversification of a protein can lead to promiscuity or pleiotropy [58, 59], especially when horizontal transfer events occur [60]. It is then likely that the protein may partially or totally lose its original function, undergoing a process of readaptation to its new genomic and environmental context [11, 61]. Such diversification constitutes the main problem when delimiting functional sequence spaces. A common method to predict function among divergent proteins is the orthology delineation, which account for speciation and duplication events, whereby their chronology allows the classification of gene sets into paralogs, orthologs, inparalogs, outparalogs, etc. [62, 63]. Yet, orthology delineation methods were less accurate than our approach in terms of functional grouping within the same protein family. Instead, our approach relies on biochemical evidence at the sequence level. We clustered the *amoA* tree by means of orthology delineation with *possvm* [64], which relies on the species overlap algorithm [65] (Fig. S8). The result was different from that obtained with our method. Some ortholog groups were correctly delineated, such as AOB (*u* = 1; OG40). However, they included methane oxidizing (*u* = 0), butane oxidizing (*u* = 0), and ammonia oxidizing (*u* = 1; AOB-gamma) sequences in the same ortholog group (OG16). Consequently, we opted for a conservative strategy, i.e., to propagate the biochemical evidence at the sequence level to robust regions of reference trees.

Once the functional sequence spaces have been properly identified, the next task is to determine what is functionally distinctive about these divergent groups. Neutral drift undoubtedly contributes to protein sub- and neo-functionalization [66]. However, the observed functional plasticity in a population of sequences may be constrained by selection, allowing different kinetics in diverse environments, even modifying the substrate preferences [67–69]. We can expect that sequence variants adapted to similar conditions will converge in similarity. Thus, if the entire sequence space performing exactly the same function is grouped into clusters of sequence similarity, groups of sequences that are expected to work alike in similar environments shall emerge (Figs. 3 and 4), instead of at-random groupings. For quantification of microbial guilds, we defined the function implementations as these groups of sequences that work equivalently towards a key resource, regardless their lineage (same binding affinity, substrate spectrum, temperature, pH or salinity preferences, etc.).

### 3.2 How the environment constrains microbial protein diversification

Like all other organisms, microbes achieve proteostasis through the expression of regulatory feedbacks, tuning of non-covalent interactions between structural subunits, and sequence re-adaptation [70–72]. All of these mechanisms act in multiple levels and can have an immediate impact on substrate accommodation [73]. A single amino acid change may be crucial for the specificity between the substrate and its binding site [74, 75]. Moreover, modification of residues at sites other than the conserved regions of the protein can often be structurally important [76]. Regarding the quaternary structure, the protein subunits evolve to remain bound under physiological conditions, and to monomerize in out-of-range environments [77]. Sometimes, due in part to the non-covalent nature of these protein-protein bonds, it is possible to recover function when physiological conditions return [77]. For all these reasons, it can be stated that any functional protein is the fine-tuned product of a sequence to a particular range of environmental conditions.

However, the process itself is poorly understood in a mechanistic way despite continued efforts [78–80]. More recently, Panja et al. statistically explored the changes in microbial proteins adapted to different types of environments, and they diverged both in terms of amino acid composition and in their arrangement [81], a result in line with the *weak selection* concept [82]. According to these and other previous results, salinity, pH and temperature would represent the major environmental drivers of how proteins evolve [4, 83–85], modifying the catalytic kinetics, substrate specificity or conditional stability while maintaining the same function [69, 86].

From these facts it is deduced that environmental constraints, beyond speciation processes and gene duplications, are key controllers of functional structure in microbiomes. Therefore, we longed to classify proteins ecologically by their ability to operate similarly accross environments.

### 3.3 Determining microbial implementations considering the nature of protein diversification

As a case study, we chose a function that is difficult to explore and quantify, which is organic nitrogen acquisition through putrescine and other related polyamines. The difficulty of exploring this function is given by the following issues: (i) substrate affinity is moderately unspecific and, although there may be a slight preferential binding to spermidine or putrescine depending on certain amino acids [87], our results indicate that proteins with the same substrate preferences are not strictly clustered in the tree (Fig. S4); (ii) there is an extreme shortage of sequences with functional experimental evidence.

As stated above, the microbial guild quantification method aims to (1) discriminate sequence spaces that correspond to the same function, and then to (2) characterize groups of functional sequences that work in a similar way (implementations of the function). The first objective improved automatic gene annotation, while the second categorized it functionally. To do both, we built several reference phylogenetic trees for oceanic organisms and used them as sequence space classifiers (Figs. 3 and 4).

Regarding objective (2), we wanted to classify the performances of ABC transporter-associated polyamine-binding proteins. Since we did not have sufficient information on the preferential binding to each polyamine or its kinetics, we decided to characterize groups that work alike in a different way. We manually obtained environmental preference information for as many of the cultured organisms present in our reference tree as possible. Then, we tested how good the tree topology was at discriminating groups of sequences putatively adapted to work in given ranges of the environmental variables.

When evaluating sequentially all nodes in the tree, we found that some internal nodes had a significant correspondence with particular environmental variables. These nodes divided the tree into large clades containing sequences that correlated with salinity, pH, and/or temperature (Fig. 4a). In addition, motility was also very significant for the same group of sequences related to pH variability. One possibility was that these environmentally consistent clades were phylogenetically equivalent. That is, the adaptation of transporter proteins to a given environmental condition should have been predicted by taxonomy. However, this was not the case. We found that the divergence of the bigger groups was not explained by taxonomy (Fig. 4b). Consequently, we defined the implementations for *polyamine uptake* as the sequence spaces corresponding to these inner nodes. Note that, until this work, the study of microbial guilds failed to consider how the functions were implemented, since organisms contributing to the guild in different ways prevented membership.

### 3.4 Decoupling taxonomy and function required a redefinition of guilds

We have argued above that taxonomic position is not, in many cases, synonymous with function. In addition to those arguments, there is some research actually focused on decoupling taxonomy from functional assets [88, 89]. Moreover, machine learning approaches appear to outperform niche prediction with functions rather than phylogeny [90]. This means that, at least in specific cases, it is possible to better predict the occurrence of function in an environment by its physicochemical features rather than by the taxonomic composition detected therein [89]. Despite this, some recent efforts have attempted to statistically infer microbial guilds through patterns of variation in species assignments and ecosystem properties related to the samples [91]. These authors were trying to find a solution without relying on automatic annotation. However, as the authors state, the taxa recovered with their model are just *likely related to a function*, but (i) we cannot know the mechanism – or how they are *related* –, and (ii) it assumes an absolute concurrence of taxonomy with function.

In this context, our guild definition has proven useful for decoupling taxonomy from how the molecular functions are carried out, and our method was robust enough to improve automatic annotation. Now, microorganisms no longer have to occupy the resource space in a *similar way* to obtain membership, but can contribute to the guild with different implementations of the function over the key resource. For example, two individuals of the same species may be exploiting the same resource at very different rates, or two individuals of different species may be equivalent in terms of consumption of the key resource; taxonomy is not an absolute predictor of either functions or how they had been implemented. Decoupling these effects was therefore mandatory to quantify microbial guilds.

As shown in the results section, the acquisition of putrescine-like polyamines is a ubiquitous trait in the examined ocean layers, which is consistent with previous literature on the topic [54]. Nevertheless, we added novel insights about how this guild changes with depth by detailing how the different species implemented the function. The guild presents itself, however, in multiple shapes; it changes both its taxonomic composition and the most abundant implementations, and seems to follow trends that correspond to the different physicochemical characteristics of the three ocean depths analyzed. In fact, there are characteristic guild patterns that seem to be better explained by depth than by sampling spot or latitude. We found that this function was carried out by a wide variety of taxa and every considered implementation of *polyamine uptake* was always present, although with different strength. This result seems to support the statement of functional redundancy being more prevalent than expected by chance in microbiomes [92].

In Bergauer’s study [54], *polyamine uptake* was one of the traits studied to analyze microbial heterotrophy in different ocean layers. Regarding the analysis of the guild, our approach adds: (i) high quality depuration of spurious metagenomic queries (Fig. 4a); (ii) discrimination of sequence spaces corresponding to a functional attribute (Fig. 5b) not deducible from taxonomy (Fig. 4b); and (iii) determination of the unexpected diversification of function’s proteins (Fig. 5c). Finally, it makes the comparison between different guilds easier, as the ecological values are standardized by the same theoretical framework.

Thus, in Figure 6, *k_meso_/k_epi_* ratio had strong responses among implementations adapted to a wide pH range (*cIIa* and *cIIb*). In our dataset, this remarkable increase is shown to be exclusive for this type of implementations, and can be explained by the rapid depth-dependent acidification, a characteristic feature of the mesopelagic oxycline [93, 94]. In general, the lowest pH levels in the water column correspond with the presence of an oxygen minimum. The decrease in pH is mainly driven by the increased concentration of dissolved carbonic acid, which also relates to biological activity of upper layers (Figure S7). Values of pH are also dependent on more strictly abiotic factors such as temperature, salinity and pressure, acting as dissociation constant modifiers [95, 96]. Therefore, the pH minimum is strongly linked to mesopelagic depths and may exhibit some seasonality. A key result was that the implementations that most changed their representation (both in abundance and richness) between epipelagic and mesopelagic were precisely those preserving their function despite pH variability. This was not only a validation of the method, but a significant step forward in knowledge of functional ecology. We can now make use of a new window to search for structure.

## 4 Conclusions

This work has three strands of contributions: (i) conceptually, we propose a new definition of guild, as previous definitions led to ambiguity, especially in microorganisms; (ii) methodologically, we have been able to formalize the new concept of guild for the study of microbiomes by delineating the sequence spaces and assigning functional properties to them, and to develop a quantification method that corrects for efficiency and expected richness of effectors, in addition to their abundance; (iii) ecologically, we have been able to quantify and describe for the first time the *polyamine uptake* guild in water columns from tropical and subtropical latitudes, identifying key players, and finding a major correspondence between certain sequence spaces of *potF* and the oceanic environment in which they perform their function.

## 5 Methods

### Construction of the marine prokaryotic genomes database (1 in Fig. S2)

To facilitate the construction of the gene-specific reference databases, we compiled a database of peptide sequences obtained from a collection of prokaryotic, quality-filtered genomes (MAGs and SAGs) from marine environments. Specifically, we retrieved genomes from the following databases: a) the MAR database, all 1,270 complete genomes, and 5,521 partial genomes that had the “high quality” status as described in [97]; b) the OceanDNA database [98], all 52,325 genomes, since they had been quality-filtered based on their completeness and degree of contamination with the formula: percent completeness - 5 × percent-contamination ≥ 50); c) the collection compiled by [99], which includes genomes from various origins such as TARA OCEANS [100] and XORG, in this case, only genomes passing the same quality filter applied in OceanDNA were kept, amounting to a total of 26,942 additional genomes. All the genomes considered had assigned taxonomy obtained with the GTDB Toolkit [101] 2.0.0 available in their corresponding databases. Finally, all sequences were merged into a single database, reads were further quality filtered with fastp 0.20.1, and sequence duplicates were removed with seqkit rmdud [102] 2.0.0 using default parameters.

### Functional marker selection (2a in Fig. S2)

The search for functional markers was carried out by means of an extensive bibliographic comparison. This methodology is based on choosing public available Hidden Markov Models (HMMs) [103] for one or several genes, trying to avoid functional paralogs to maximize functional univocity. In order to choose an HMM as a guild marker, we followed the following conservative criteria: (i) the construction of the HMM must be congruent with the sequences that have reviewed functional evidence in literature, (ii) the metagemonic sequences retrieved with the tested HMM can be filtered out by a specific quality argument, derived from the inner workings of genomic architecture (i.e.: synteny) or a consequence of the evolutive history of the gene (i.e.: similar sequences that have undergone functional drift). With this methodology, we selected the best minimal markers for the guilds analyzed in this work.

### Construction of the gene-specific reference database (2b in Fig. S2)

We used the selected profile HMMs and HMMER3 [104] to retrieve candidate sequences of the target gene from our collected marine peptide database. Gather score thresholds were used as a quality filter when available, otherwise, a minimum E-score threshold of 10^−9^ was employed.

To facilitate inference and later visual inspection of the phylogenetic trees, sequence hits were further filtered to set a maximum database size of N representative sequences. To this end, we applied a series of filters. First, we set minimal and maximal sequence length cutoff values of *l1* and *l2*, respectively. Second, we removed sequence duplicates through seqkit’s rmdup sub-command with default parameters. Third, we applied CD-HIT [105] with default parameters to reduce redundancy in the peptide database. Finally, if the database size was larger than the allowed maximum after applying CD-HIT, we further reduced the number of representative peptides through RepSet [106], an optimization-based algorithm that obtains a series of nested sets of representative peptides of decreasing size. Specifically, we selected the maximal set of representative peptides with a size lower than the established size threshold value.

### Usage of synteny during gene-specific reference database construction (2c in Fig. S2)

In some cases, we used syntenic information to reduce uncertainty due to the potential presence of paralogs during the reference peptide database construction. To this end, we employed the Python package Pynteny [107], which facilitates synteny-aware profile, HMM-based searchers. After generating a list of synteny-complaint target-peptide matches, we followed the same protocol to reduce database size when required to meet the established reference database size threshold value.

### Inference of the gene-specific reference trees (3a in Fig. S2)

Once the peptide reference database was constructed, we employed MUSCLE [108] with default parameter values to perform a multiple sequence alignment of the reference database. Next, we used the previous alignment and IQ-TREE [109] with default parameter values to build a reference phylogenetic tree for each target gene. We determined the substitution model through ModelTest [110] by selecting the model with the highest AIC score.

### Classification of clusters within the reference phylogenetic tree (3b in Fig. S2)

Once we have constructed the reference tree, we can now propagate functional information that corresponds to the different regions of the reference tree. In the case of *amoA*, the functional information was obtained directly from the sequences we used to build the tree, and the clusters inferred from the similarity between sequences.

In the case of polyamine binding reference tree, we needed other criteria to classify clusters. To check whether different clusters are associated with different environmental conditions, we carried out an extensive literature search of the environmental preferences of 321 species that matched 478 leaves (41%). We assembled a curated collection of physicochemical preferences for these species that included tolerance ranges and optimal values of temperature, salinity and pH, as well as other variables such as motility (Table S1). For each internal node we calculated the average values of all its leaves. To determine whether the association with environmental variables of the cluster were significant, these node averages were compared to the distribution observed under 2·10^4^ randomizations to obtain their z-scores. Nodes with an average value of the z-score larger than 3, i.e. p-value ≤ 0.003 were considered significant for the particular environmental variable (Table S2). To select the most general internal nodes, we focused on those that are significant but whose parent node is not. These are color coded in Figure 4, 5 and 6.

To determine whether these sequence clusters were expected by the taxonomy of the organisms, we constructed a null model of phylogenetic divergence with two phylomarkers (16S and *rplB*). We then compared the divergence of *potF*-like sequences for the same organisms, finding that taxonomy does not explain, in most cases, the drift found in functional genes; however, environmental variables do, especially for nodes that separate more leaves on the tree, as shown in Figure 4b.

This methodology can be applied to any type of functional evidence, not only environmental, but also kinetic, substrate preference, or any other type.

### Preprocessing of query sequences (4a in Fig. S2)

Query sequences were retrieved from the metagenomes following SqueezeMeta’s pipeline [111].

### Placement of query sequences (4b in Fig. S2)

To place query sequences (metagenomic output) in the reference tree, we first obtained an alignment between the query and the reference sequences with PaPaRa [112] 2.5. Then, we placed query sequences with the tool EPA-ng [113] 0.3.8. Additionally, we employed the Gappa toolkit 0.8, specifically, the command gappa examine graft [114] to visualize the placed sequences on the reference tree using default parameters. The phylogenetic placement tree was visualized using the Interactive Tree of Life [115].

### Taxonomical and functional labeling of placed query sequences (4c in Fig. S2)

We employed the Gappa toolkit [114], specifically, the command *gappa examine assign* to assign taxonomy to placed sequences. Briefly, Gappa first assigns a consensus taxonomy to each internal node of the tree and then assigns to each query sequence the closest taxonomy in the reference tree weighted by the likelihood of each placement. We employed default parameters and the *best_hit* option to retrieve only the taxonomic assignments with the highest total placement likelihood for each query. To assign functional labels to placed queries, we selected the function of the tree cluster in which each query had been placed. To this end, we first added the cluster label to each taxonomic path of the reference sequences as an additional (artificial) taxon above the domain level. In this manner, we could employ *gappa examine assign* to assign both taxonomy and the cluster label (i.e., function) to each placed query.

### Quantification of the *polyamine-uptake* guild from metagenomic data (5 in Fig. S2)

Further technical details regarding the application of our method can be found in the Supplementary Information and the repository tutorials (https://github.com/pyubero/microguilds/tree/main/Tutorials).

### Expected sequence richness model

In our data, we invariably observed that larger sequence abundance corresponds to larger sequence richness, this arises naturally when sampling a large pool of distintct elements far from saturation. Briefly, we assess the expected richness by fitting

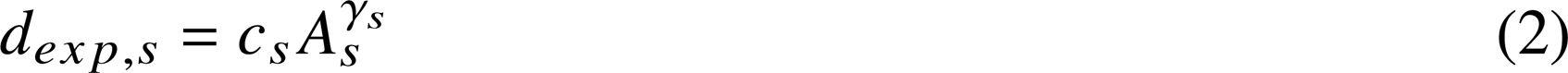

where *s* are the sequences considered for a taxon, implementation and environment, abundance is measured in TPM, richness is the natural number of unique sequences, and where *c*_s_ and *γ*_s_ are constants. We consistently obtained *R*^2^ > 0.8 (Fig. S9). Consequently, when computing the impact coefficients in Eq. 1, the term *d_obs,s_/d_exp,s_* rewards sequences that display a larger richness than expected.

### 16S and rplB sequences

To screen phylogenetic deviations between functions and phylomarkers (Fig. 4b), we obtained the nucleotide sequences of the 16S ribosomal subunit and the rplB gene from the assembly genomic RNA and CDS provided by the NCBI for 319 out of the 321 species found in pure culture. When 16S sequences were <1000bp, we used instead sequences from other strains as they should remain well conserved within the same species. All RefSeq assembly accession numbers and alternative GIs for 16S data were automatically retrieved from the NCBI, a detailed list is available in the Supplementary Table 4.

## 6 Akcnowledgements

We are grateful to Juan F. Poyatos (CNB-CSIC) for discussion of the methodologies and their implications, to Alberto Pascual-García (CNB-CSIC) for the feedback and discussion on the epistemology of the microbial guild, and to Diego Jiménez for encouraging this work. We also thank the CSIC-LifeHUB forum (PIE-202120E047-Conexiones-Life) for generating space for discussion and allowing these ideas to mature.

## 7 Author contributions

J.R.S.: Conceptualization, Methodology, Validation, Writing - Original Draft, Investigation, Data curation, Visualization; P.Y.: Methodology, Software, Formal analysis, Writing - Review & Editing, Data curation, Visualization; S.R.E.: Methodology, Writing - Review & Editing; J.M.G.: Methodology; J.T. and C.P.A.: Writing - Review & Editing, Project administration, Supervision, Funding acquisition.

## 10 Materials and correspondence

Correspondence and request for materials should be addressed to J.R.S.

## 11 Data availability statement

All data generated or analysed during this study are included in this published article and its supplementary information files.

