## Supplementary material for "Quantifying Microbial Guilds": Supplementary.pdf

### Title: Quantifying microbial guilds

---

Supplementary Information

#### Discussion: Determining microbial implementations considering the nature of protein diversification

In addition to the things explained in the main text, we reported that some terminal nodes were enriched in significance by the same or other variables. The significant environmental variables were more related to taxonomic position and vertical inheritance it the nodes closest to the leaves (Fig. S5). Some of these smaller groups contributed to the significance of a variable for a corresponding parental node. Others were small clusters, that might be caused by a lack of data or because it is a highly conserved property around a genus or species. This is the case, for example, with the ability to grow on aromatic hydrocarbons. This variable (PAHs presence in the environment) was found in a multitude of small monophyletic groups, but we never found a cluster in the tree that was significant for this variable. In addition, when we computed phylogenetic distances between representatives from these PAHs-significant groups, we noticed an increased divergence in *potF*-like sequences (0.262 normalized phylogenetic distance between *Marinobacter lutimaris* and *M. litorale*) compared to their 16S rRNA sequences (0.029). When they were organisms belonging to the same species (~ 0.0 of divergence in 16S), their divergence in *potF*-like sequences was highly variable. When comparing representatives from different genera, we still observed the increase in divergence. For instance, the divergence between *Pseudomonas alcaligenes* and *Neptunomonas antarctica* in their 16S rRNA sequences is 0.195 and 0.337 in *potF*-like sequences. Thus, we observed that organisms with this property tend to be more related in taxonomic position (inferred by a phylomarker) than in their polyamine-binding proteins. With this evidence, we concluded that PAHs presence is not an environmental feature strongly compromising polyamine acquisition effectors, and the changes in the significant sequences are better explained by other variables and taxonomic position.

#### Discussion: Unexpected richness of an implementation justification

As explained, our model rewards versatile behaviors for the same function in an environment, inferred by the richness of its effectors, as long as $d_{obs}$ is greater than $d_{exp}$. This is because we postulate that unexpected sequence diversification increases the odds that the function will persist in the environment when exposed to undefined changes. This phenomenon, although not formally described, has been hinted at in a multitude of different biological systems (Wright, 2005; Hakes, 2007; Fohse, 2011; Soyer, 2010).

This correction was introduced to estimate the importance of a molecular function because of the following reasons: (i) empirically, we observed that each gene grows in richness of unique sequences differently with relative abundance, as seen in Figure S8; (ii) abundance values for lower than expected richness can be explained by the strong dominance of an organism in a particular sample, but it does not imply that this function is responsible for the ecological success of the dominant organism, so low-richness abundances will be overestimating the importance of the function; (iii) higher than expected richness should result in a higher function robustness, since the loss of fitness for the global function regarding environmental changes should be reduced as the sequence space widens.

#### Methods: Quantification of the *polyamine-uptakers* guild

Once filtered sequences are classified by environment, taxon, and implementation, they are merged together with the corresponding normalized abundances into a single master table. This is the input for the first tool of our public repository, a python module called *guild_tensor_generate.py*.

The module will extract all the required information for the calculation of each implementation, taxon and environment-dependent functional contribution, *k*. In the present case, we study three distinct environments, so the software will produce an array of dimension 3 x *m* x *l* (where *m* is the number of implementations established within the guild marker, and *l* the number of taxa).

The calculation contemplates three terms. The abundance, A_{s}, has been calculated as a summation of normalized metagenomic counts for all the sequences contained in the same implementation, taxon, and environment. The second term is d. Theoretical d is the unexpected sequence diversification according to the sum of A_{s}, the first term. Calculation of the theoretical d is complex and would require avoiding false negatives. Therefore, in our work it is limited by our ability to retrieve this kind of data. Finally, the term representing the univocity of the implementations, u, is equal to 1.0 since we discard the metagenomic sequences falling into non-functional sequence spaces of the reference tree, or false positives. So, our ability to determine the functional sequences is dependent of the experimental evidence for specific sequences.

In addition, we had a highly-conservative criteria to estimate the functional sequence space, as described in Methods. Ideally, environmental inhibition of the effector must be considered for the univocity calculation, but since we lack the data, we have decided that there is no inhibition for this example. An example of the k-tensor output is provided in the Supplementary Table 3.

The second tool, *guild_tensor_visualize.py,* helps to visualize this tensor, which can be of varying complexity. It does two things: (i) it filters by the taxonomic level to visualize the guild patterns and (ii) it takes the value of *k* by taxonomic contribution to each implementation and environment. To do the latter, it takes the contribution of each position in the tensor and plot them with different preferences top contributors or rare taxa, linear or log representation, polar or rectilinear charting, etc. as shown in Fig. 5, resulting in an easy way to visualize complex data.

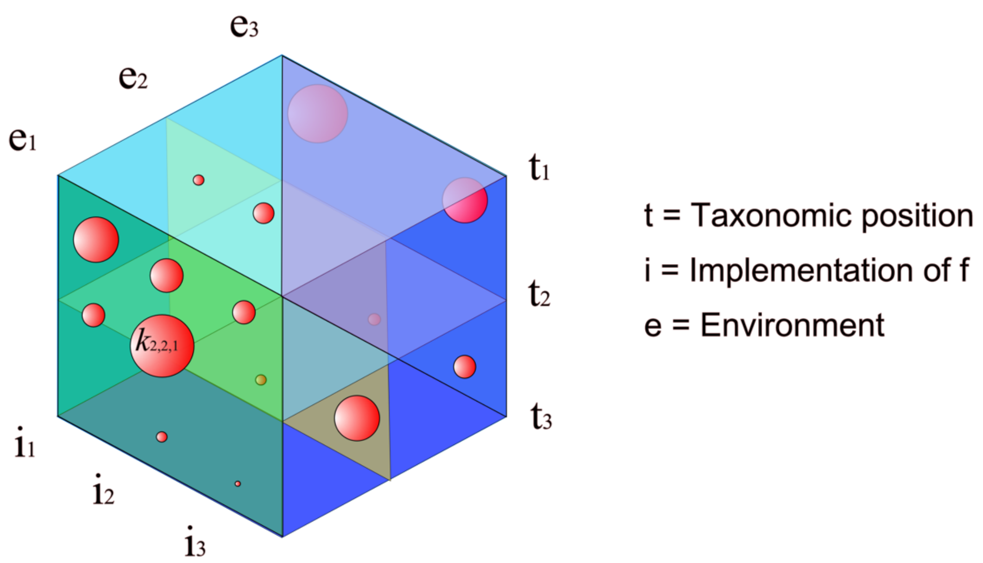


**Figure S1 – The ecological dimensions of a guild.** Visual concept of the three-dimensional matrix we used during the quantification of the contribution to the guild in different ecological dimensions. The guild structure can be defined as each of the impact coefficients *k* that the definitory function has on each triplet taxon, implementation, and environment.

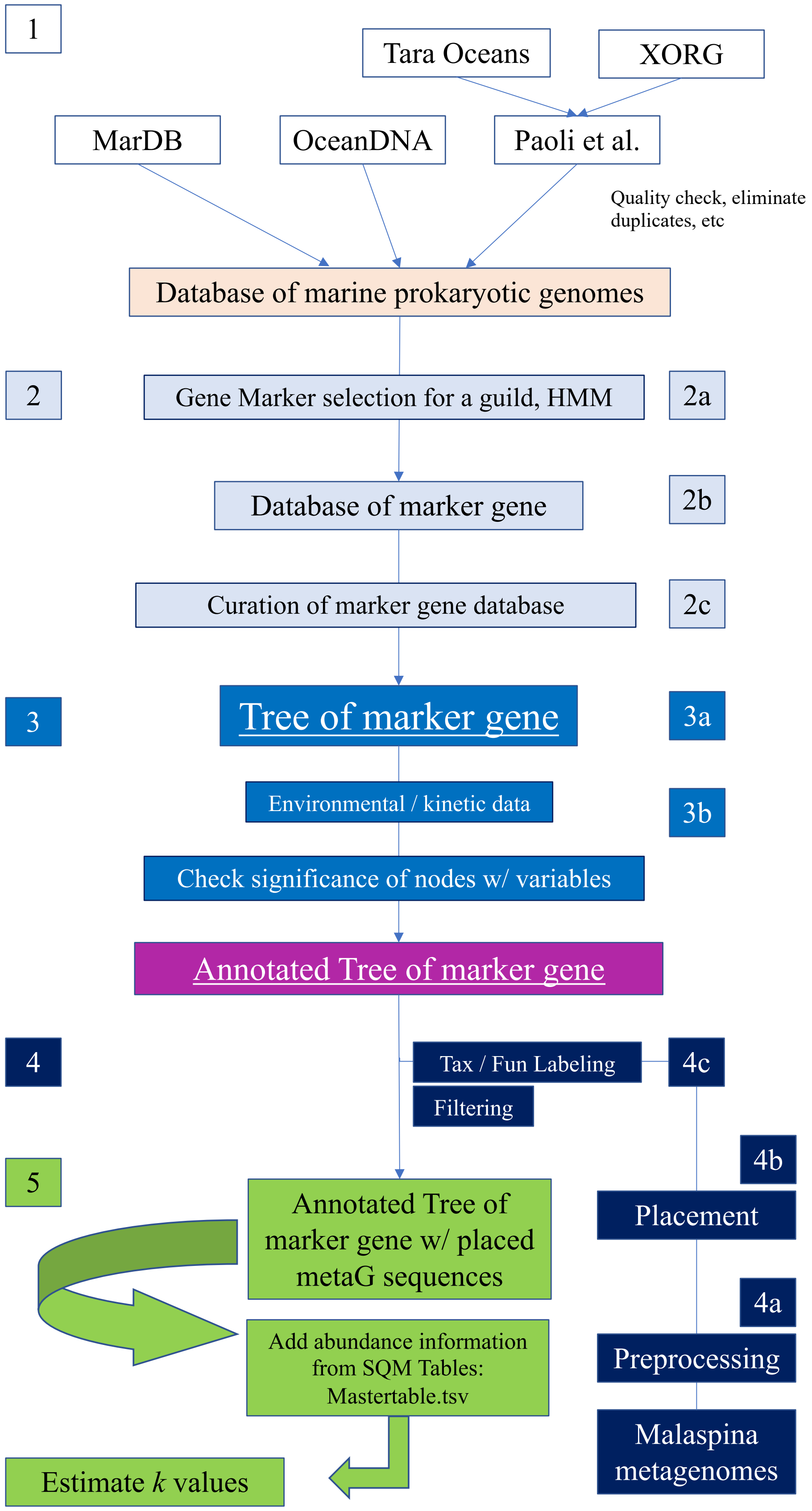


**Figure S2 – Complete microbial guild assessment workflow.**

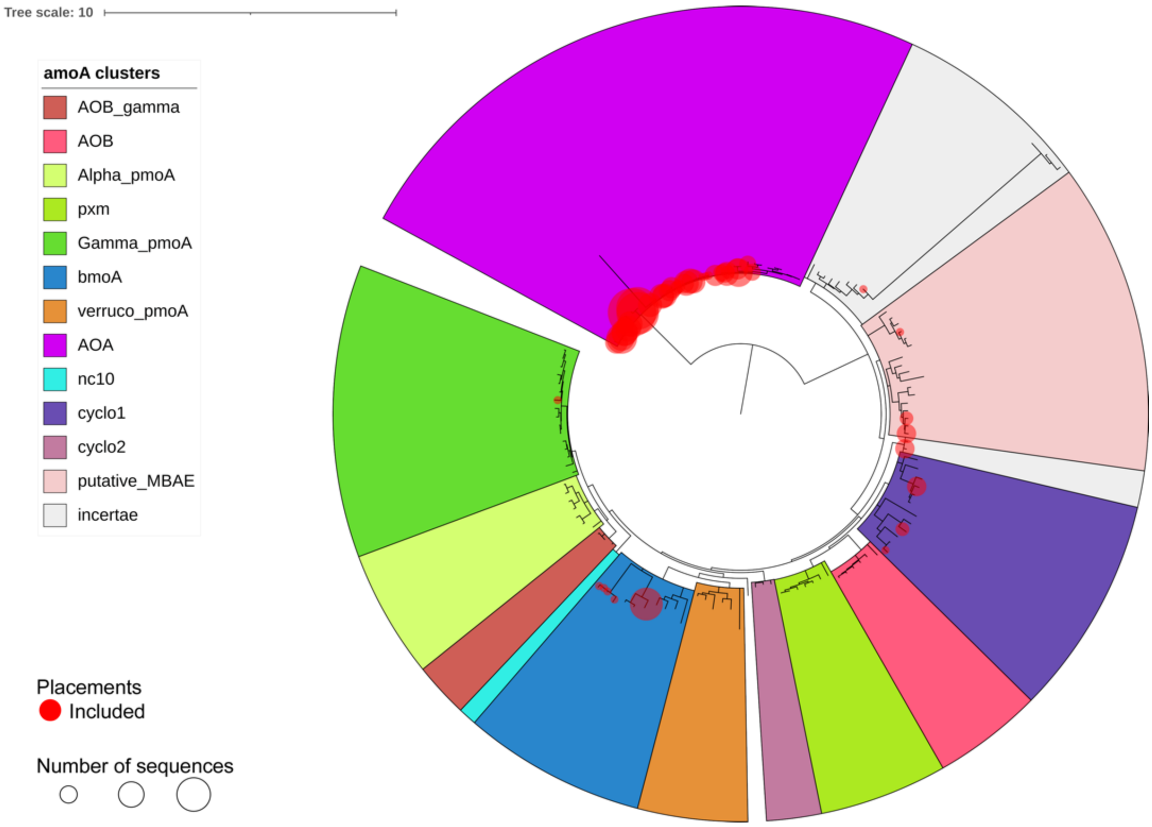


**Figure S3 - Reference tree and metagenomic placement (red dots) for *amoA*-like sequences, used as a marker for ammonia oxidation function in the ocean.** Clusters labeled AOB and AOA are truly ammonia oxidizing sequences. The rest are known non-univocal sequences, oxidizing methane (*pmoA*) or carrying out other functions. Most environmental sequences were AOA, showing the importance of archaea in the open ocean. There were also sequences in some of the non-univocal clusters. These can be either discarded or retained for comparison. In either case, the number of sequences assigned to ammonia oxidation is not artificially overestimated with this method.

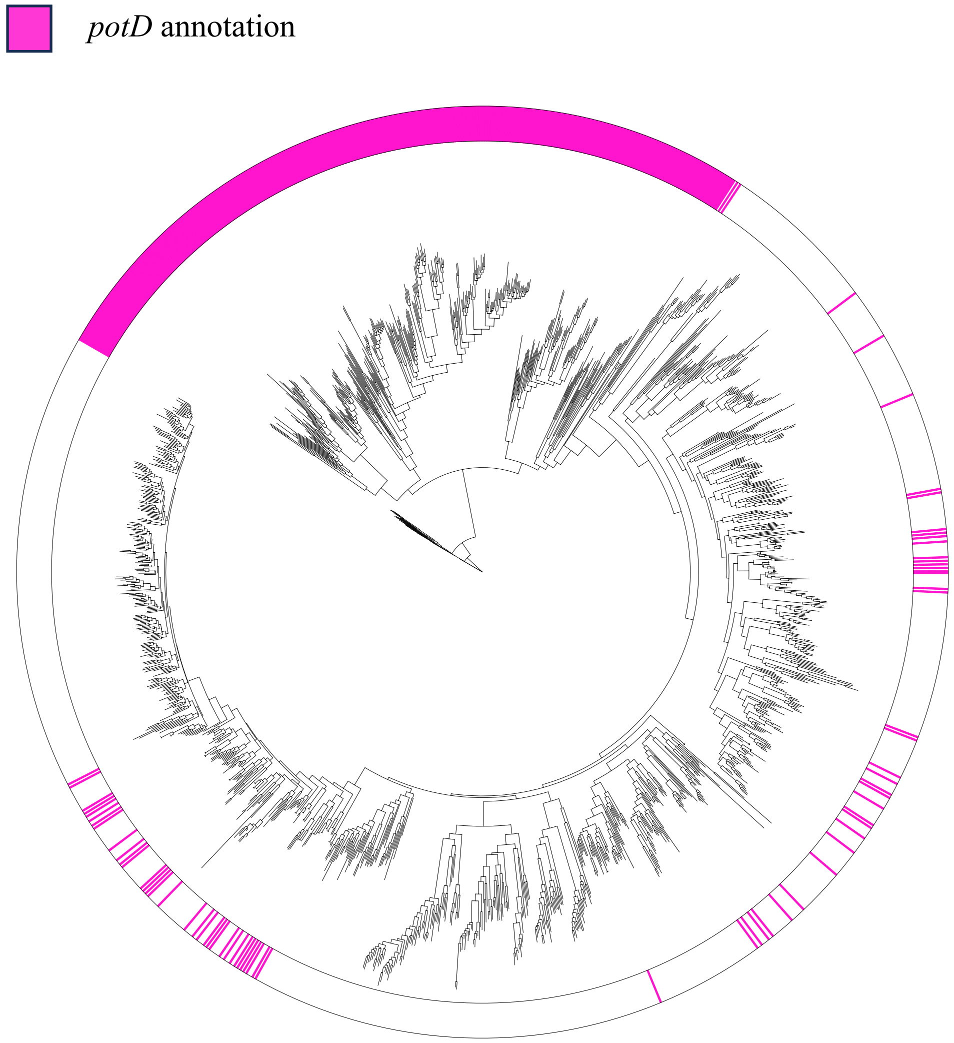


**Figure S4 - Phylogenetic tree considering the notation of two different genes, *potD* (pink) and *potF* (white).** *potD* is an annotation corresponding to a *potF* ortholog. Affinities for spermidine and putrescine are slightly different between *potD*/*potF* annotations. As stated in the main text, we wanted to know whether we could distinguish precisely between sequence spaces with a preference for binding to spermidine or putrescine. For this purpose, we reconstructed a phylogeny with the potF-like reference tree sequences and the first five hundred best matches of *potD*-like annotated sequences (retrieved with BLAST). As can be seen in the image, the branch fully annotated as *potD* is not ambiguous for annotation, while the opposite is not true for the *potF* region. This, in addition to the great substrate spectrum that these ABC transporter subunits have (putrescine, spermidine, cadaverine and other polyamines) makes hard, if not unfeasible, to determine implementations with preferential binding for this function. Therefore, we decided not to use substrate preference as a criterion for defining the implementations.

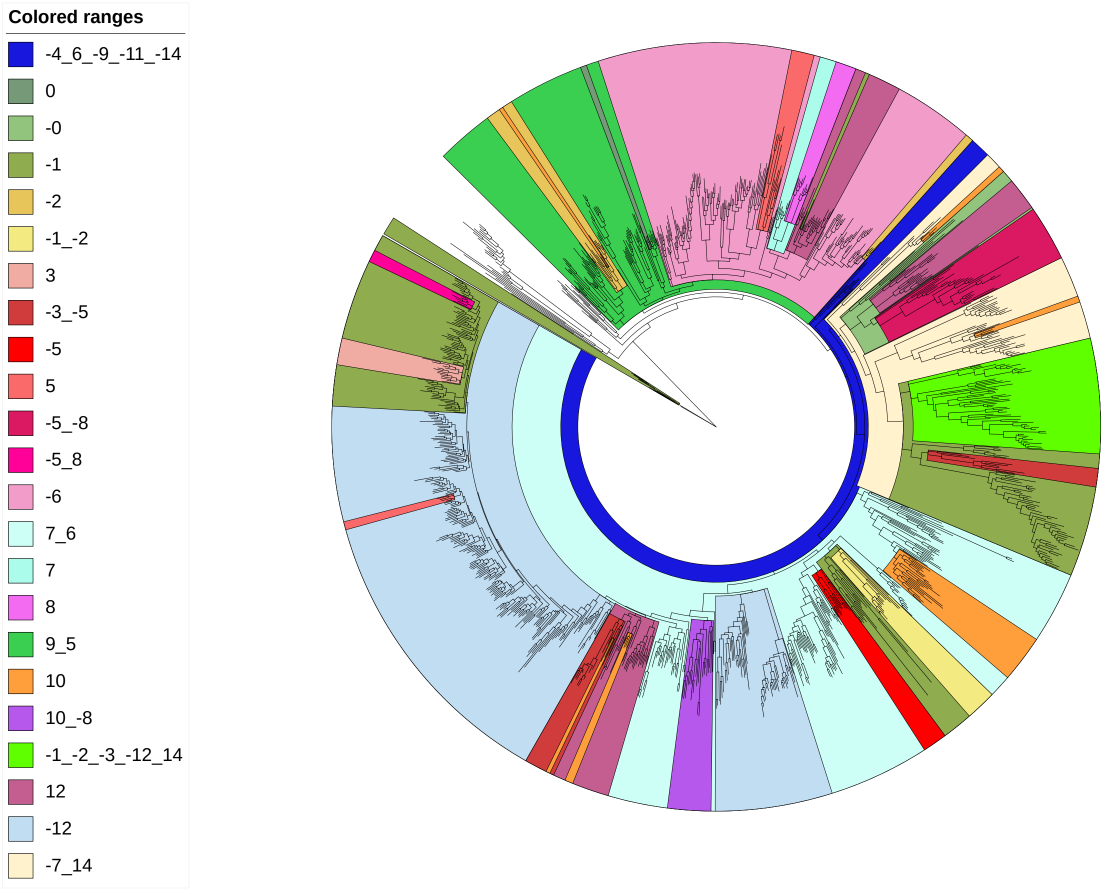


**Figure S5 - Reference tree for polyamine uptake marker gene (*potF*-like) with all significant nodes for all environmental variables tested with z-score > |3|.** The different colors indicate clusters with significant associations with environmental variables. The numbers to the right indicate whether the association is positive or negative and the number indicates which variables are significant for each clade: 0, minimal temperature; 1, maximal T; 2, optimal T; 3, minimal pH, 4, maximal pH; 5, optimal pH; 6, minimal salinity; 7, maximal salinity; 8, optimal salinity; 9, motility; 10, presence of hydrocarbons in the environment; 11, aerobiosis; 12, anaerobiosis, 13, Gram stain; 14, positive nitrate-reduction test. The outermost nodes are significant for environmental variables more linked to vertical inheritance. For instance, the presence of hydrocarbons (feature number 10, in orange) is a divergent property for this phylogenetic reconstruction. This can be explained by the fact that bacteria need a lot of functional elements for degradation. These external nodes are therefore likely to be widely distributed in trees like this one, which inaccurately captures the real phylogeny (as demonstrated in Fig. 4b). On the other hand, in a tree reflecting real phylogenetic relationships, these little groups should ideally be closer.

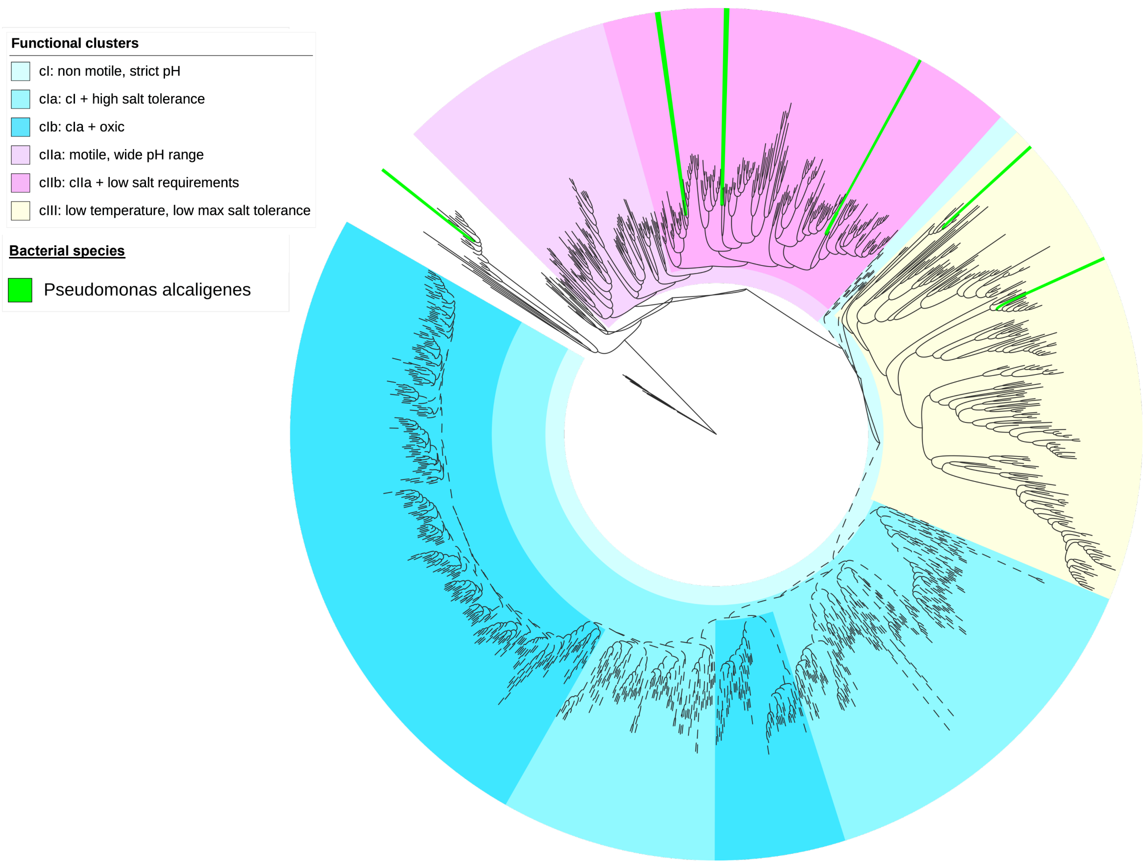


**Figure S6 – Sequence dispersion of *Pseudomonas alcaligenes* over the reference tree for polyamine uptake marker gene (*potF*-like).** We note that the same species can have phylogenetically very distant protein sequences. In the case of *Pseudomonas alcaligenes,* we found many different sequences. Some were similar to each other, but other were divergent, as shown above. *Pseudomonas alcaligenes* can hold *potF*-like sequences from, at least, three different clusters. The colored clusters have a significant relationship with one or more environmental variables. Thus, cIII includes bacteria with the widest range of temperatures for growth, but it is also within cluster I that includes non-motile bacteria with a narrow range of pH values for growth.

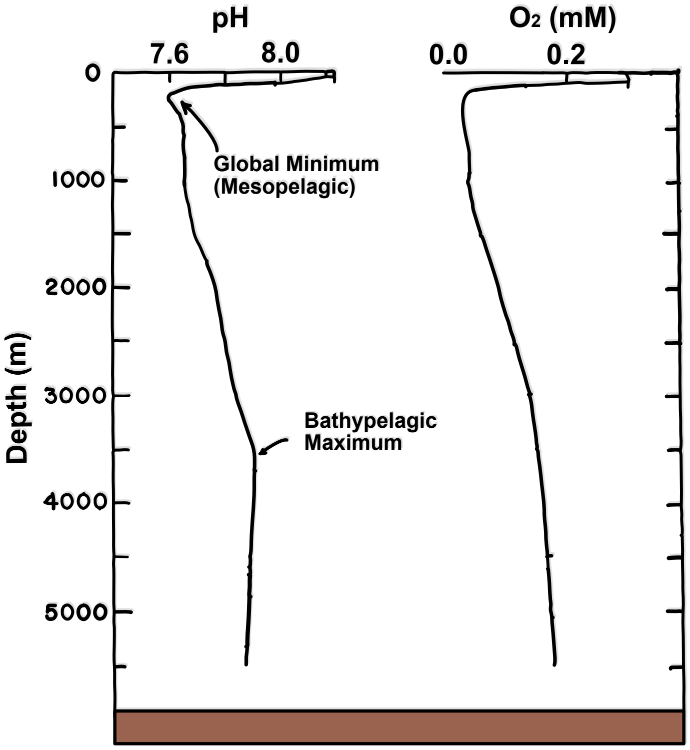


**Figure S7 – Common pH variation with depth and Oxygen concentrations.** Representative vertical profile of pH variation with depth in the ocean. (Modified from Park, 1966). Note that the mesopelagic (200 to 1000 m) experiences the largest variations in pH values. Thus, it makes sense that the most remarkable changes in polyamine binding implementations in the mesopelagic correspond to those implementations that were also significant for a large pH range.

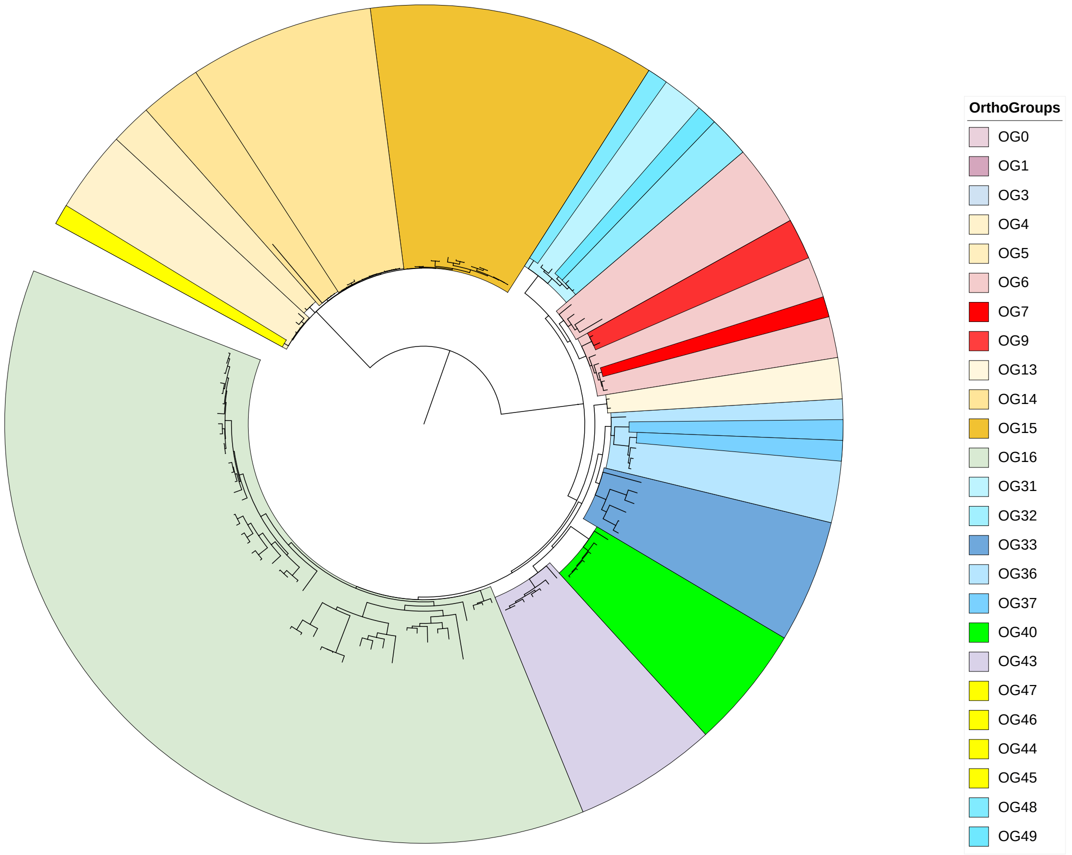


**Figure S8 – Clustering of the reference tree of *amo*A via orthology delineation failed to assess true functional sequence groups.** Although some groups were correctly delineated (OG40, corresponding to AOB amoA sequences), OG16 involved sequences with known biochemical evidence for butane, ammonia, and methane oxidation, interchangeably. Likewise, the archaeal amoA was clustered in a different way, when there is experimental evidence that all archaeal sequences belonging to AOA behave in a similar way, without promiscuity for other substrates, contrary to AOB. Therefore, we determined that using orthology-based clustering to predict function is not always a viable option, as in this case. Compare this clustering with the one provided in Figure S3.

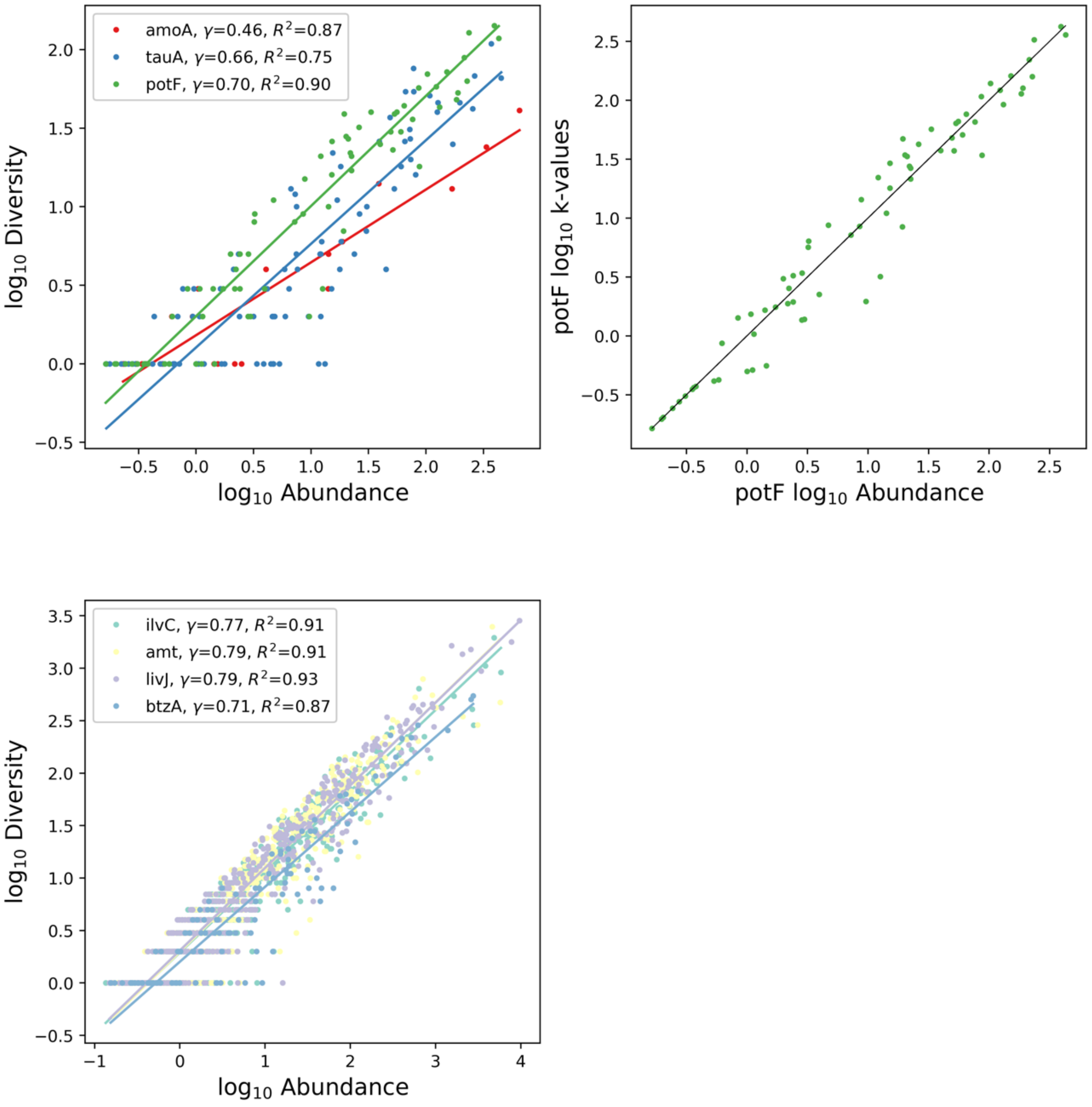


**Figure S9 – Biological data showing the empirical relation found between abundance and richness of protein sequences.** Different genes, environments and taxa show different slopes and intercepts, which are constant and allow us to calculate the expected richness in each case. Above, we have collapsed the taxonomic level to Family, while below the regression is at the Species level. Although the number of points is lower for Family, the log-log relationship holds, allowing us to anticipate several unique sequences expected for an observed level of abundance.

Supplementary Tables

#### Table S1

"Sup_table_1_environmental_data.csv" – Manually-compiled table showing the environmental preferences of each of the organisms grown and described from the reference tree for *potF*-like sequences.

For each microorganisms the following data are indicated: name, minimal, maximal and optimal temperature for growth, minimal, maximal and optimal pH, minimal, maximal and optimal salinity, motility, growth with hydrocarbons, aerobiosis, anaerobiosis, Gram stain, nitrate-red, and the literature source of this information.

#### Table S2

"Sup_table_2_A_zscore3_significant_nodes_potF.tsv" - Table where, after twenty thousand randomizations, significance values for nodes exceeding a minimum z-score of |3| (one-tailed p-values of 0.003) are observed.

Positive values indicate that the node is significantly above the mean and negative values that it is below the mean. Thus, for example, node 1 has a z score of -0.38 for minimal temperature. Although it is below the mean it is not significant. On the other hand, this node is significant for maximal temperature with a score of -3.78.

#### Table S3

"Sup_table_3_kmatrix.tsv" – Example of guild object as generated by our tool (https://github.com/pyubero/microguilds).

The table shows for each microorganisms having the potF gene the name, context (in our dataset, depth layer in the ocean), the cluster where it appears in the tree, its abundance (number of identical reads in metagenomes), diversity (number of unique sequences in the cluster), univocity (here considered to be maximal, 1), delta value (considering expected diversity and real diversity) and calculated *k* value.

#### Table S4

"Sup_Table_4_table_organisms.xlsx" – List of organisms and accession numbers for the organisms present in Sup. Table 1.

Supplementary Files

#### File S1

"FileS1-ref_database_relabel.newick" – reference tree for amoA sequences
"FileS1-mp_query_amoa" – Malaspina metagenomic query for amoA-like sequences
"FileS1-arqueabactammoa.hmm" – Built model for amoA-like proteins

#### File S2

"FileS2-tree_potF_labelled.newick" – reference tree for potF sequences.
"FileS2-putrescine_transport.fasta" - Malaspina metagenomic query for potF-like sequences.
"FileS2-K11073.hmm" – Used model for potF-like proteins
