## Supplementary figures and images for "Quantifying Microbial Guilds"

### Figure_S1.png

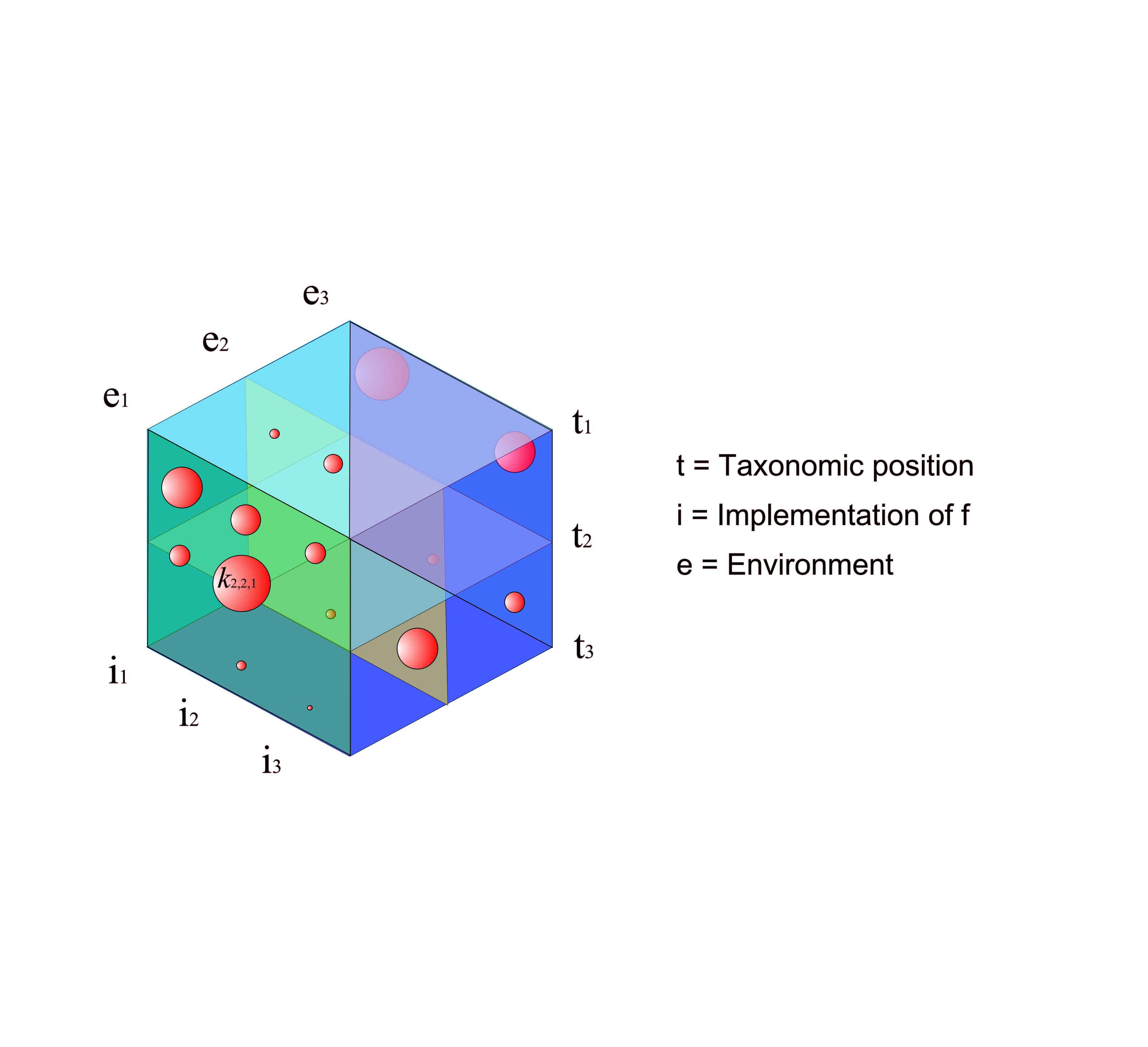

### Figure_S2.pdf

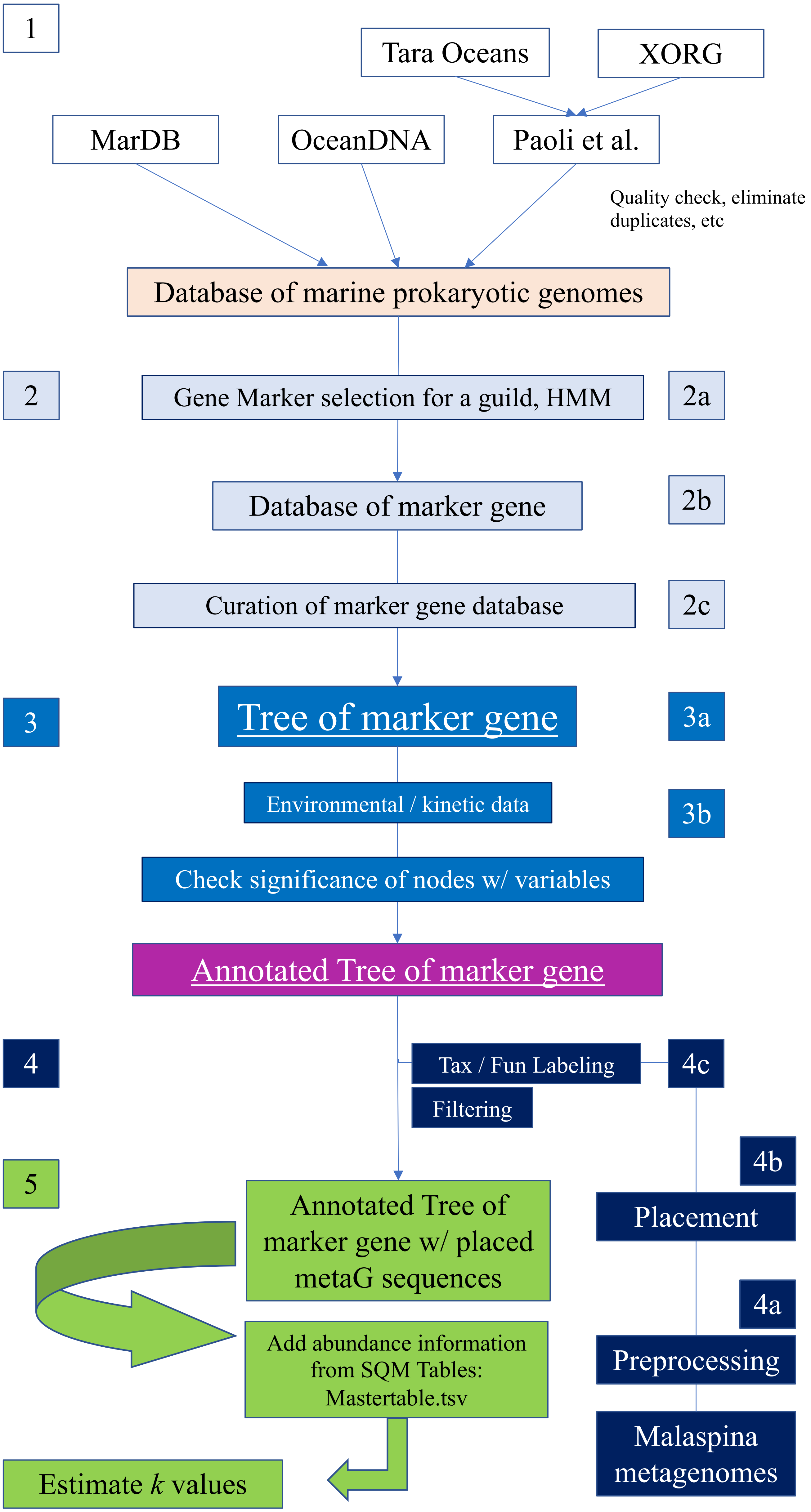

### Figure_S3.png

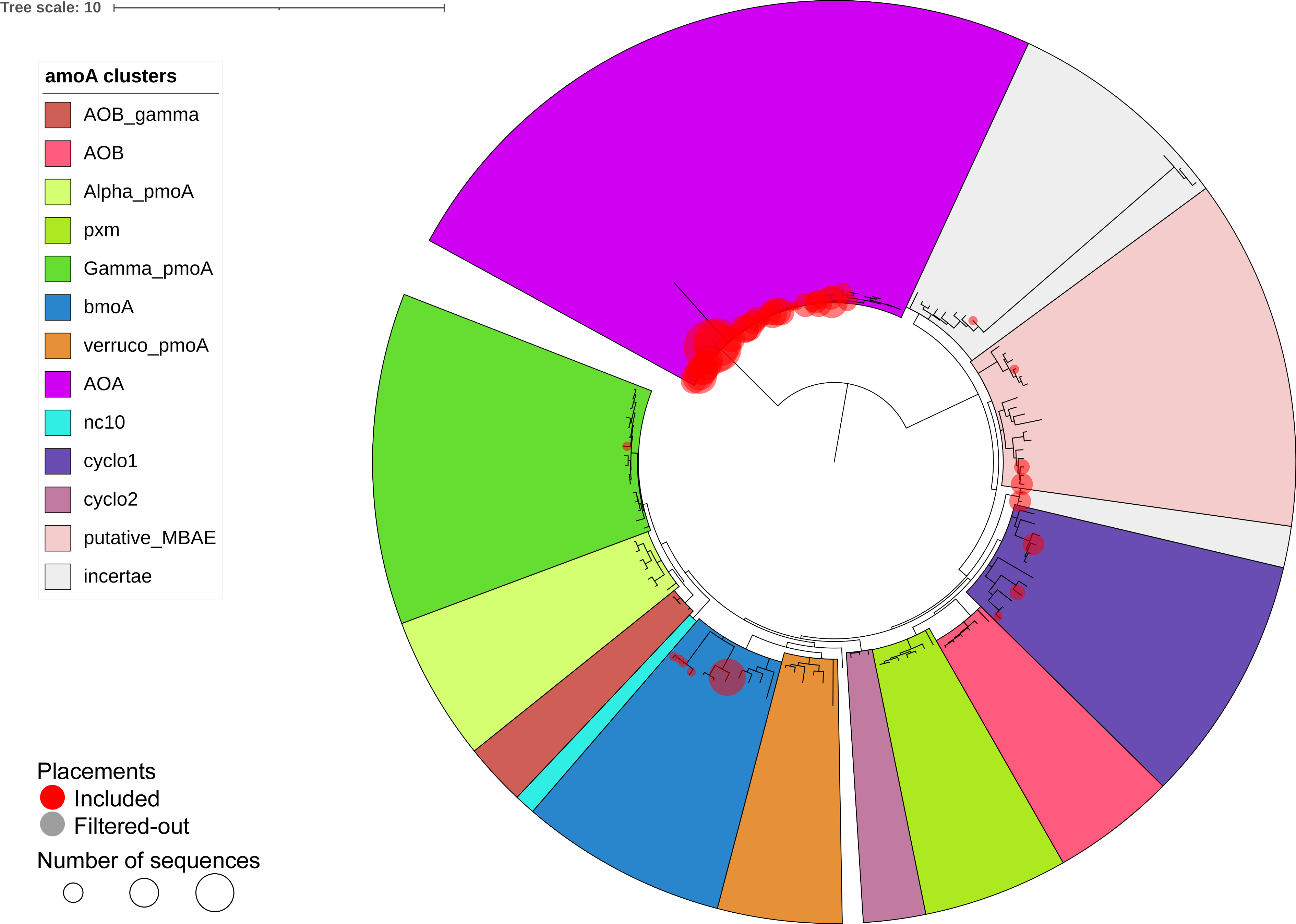

### Figure_S4.pdf

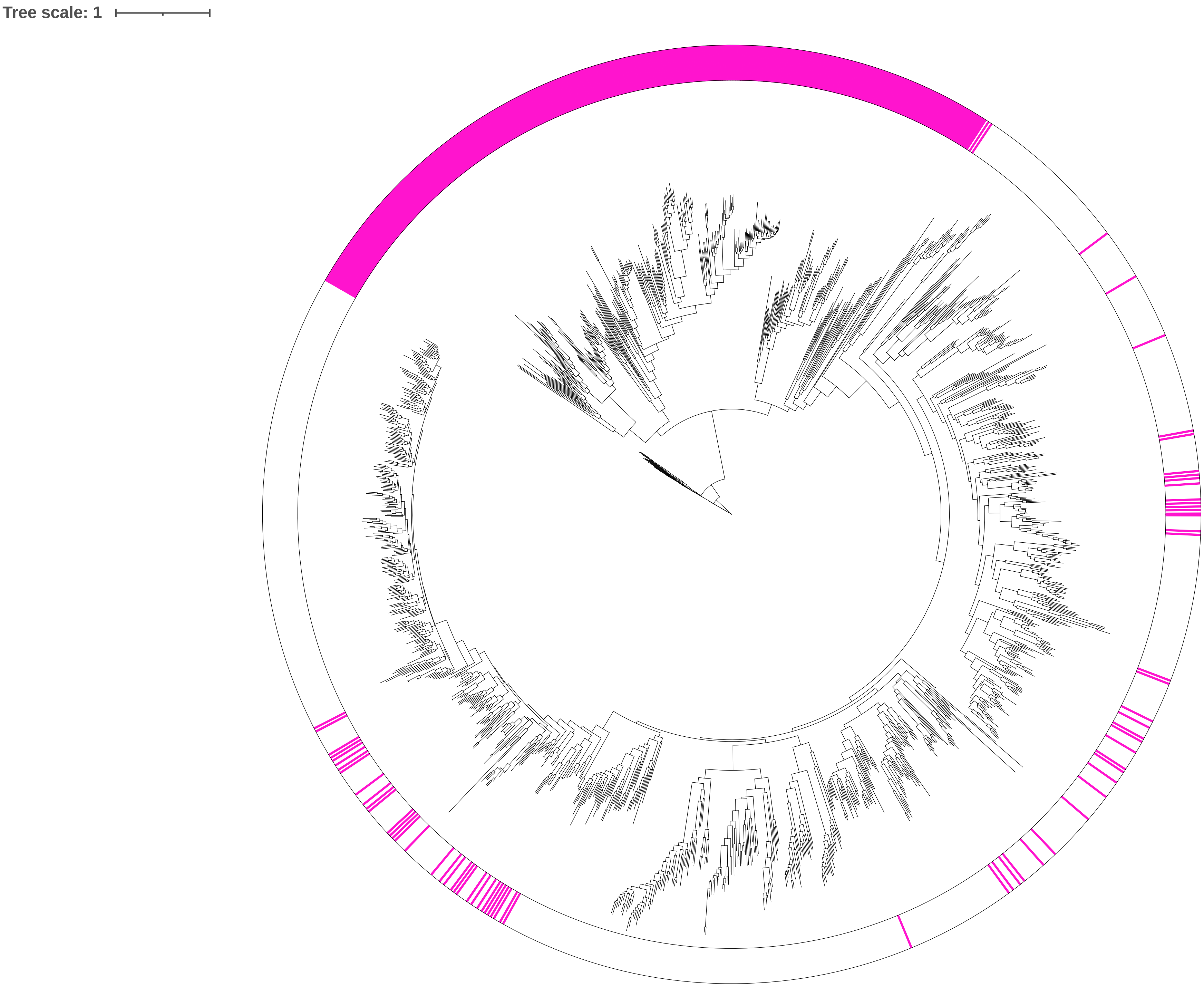

### Figure_S5.pdf

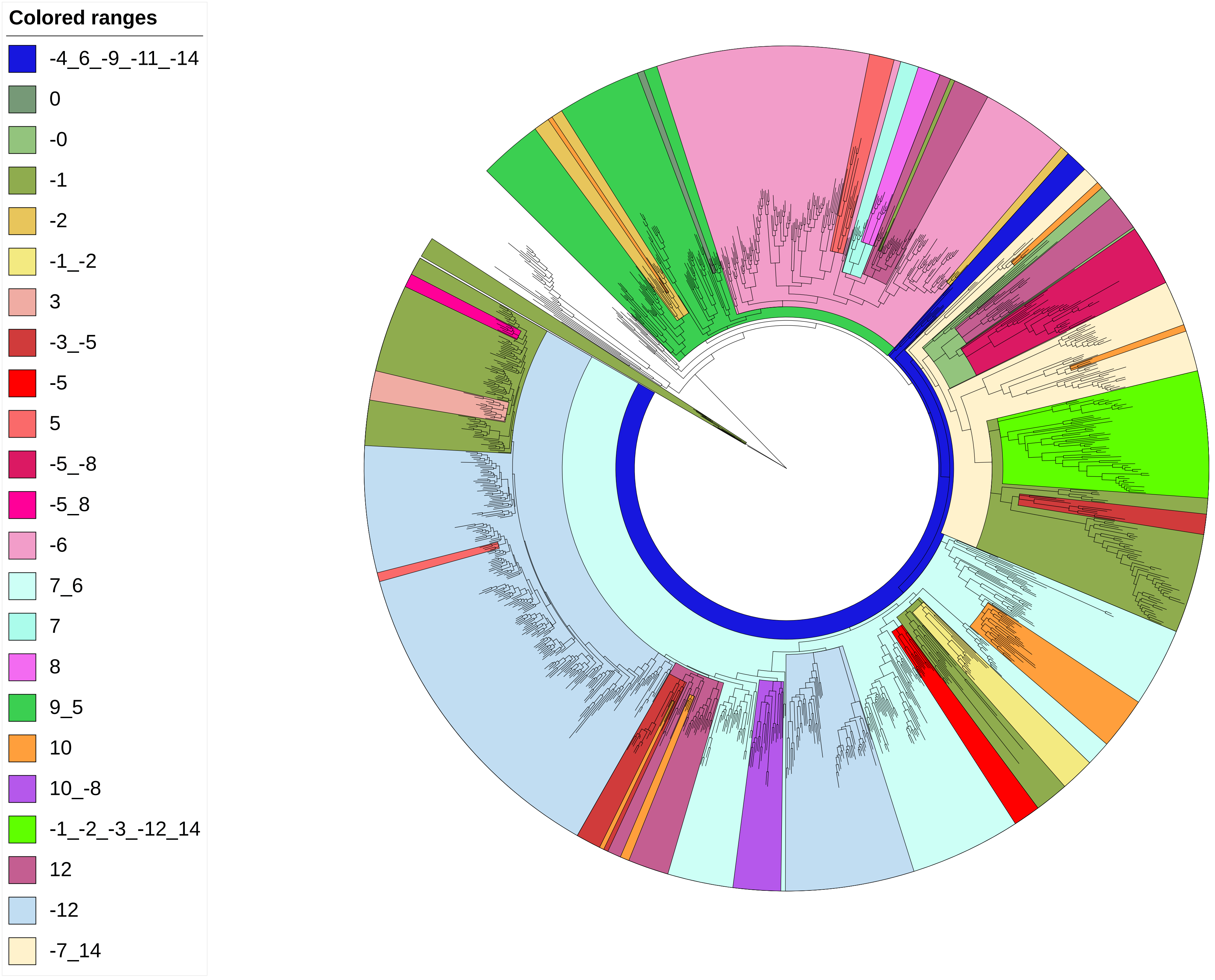

### Figure_S6.pdf

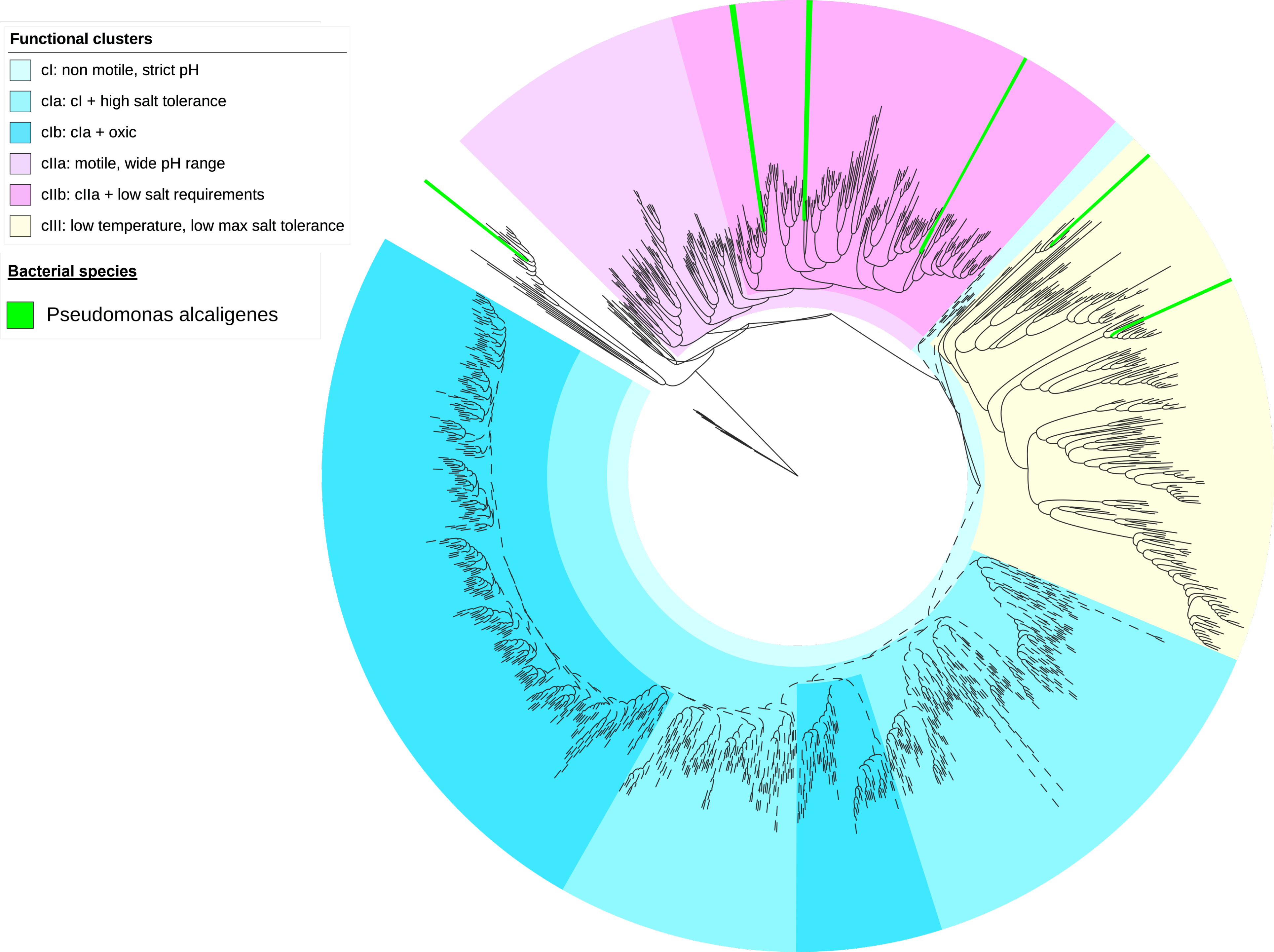

### Figure_S7.pdf

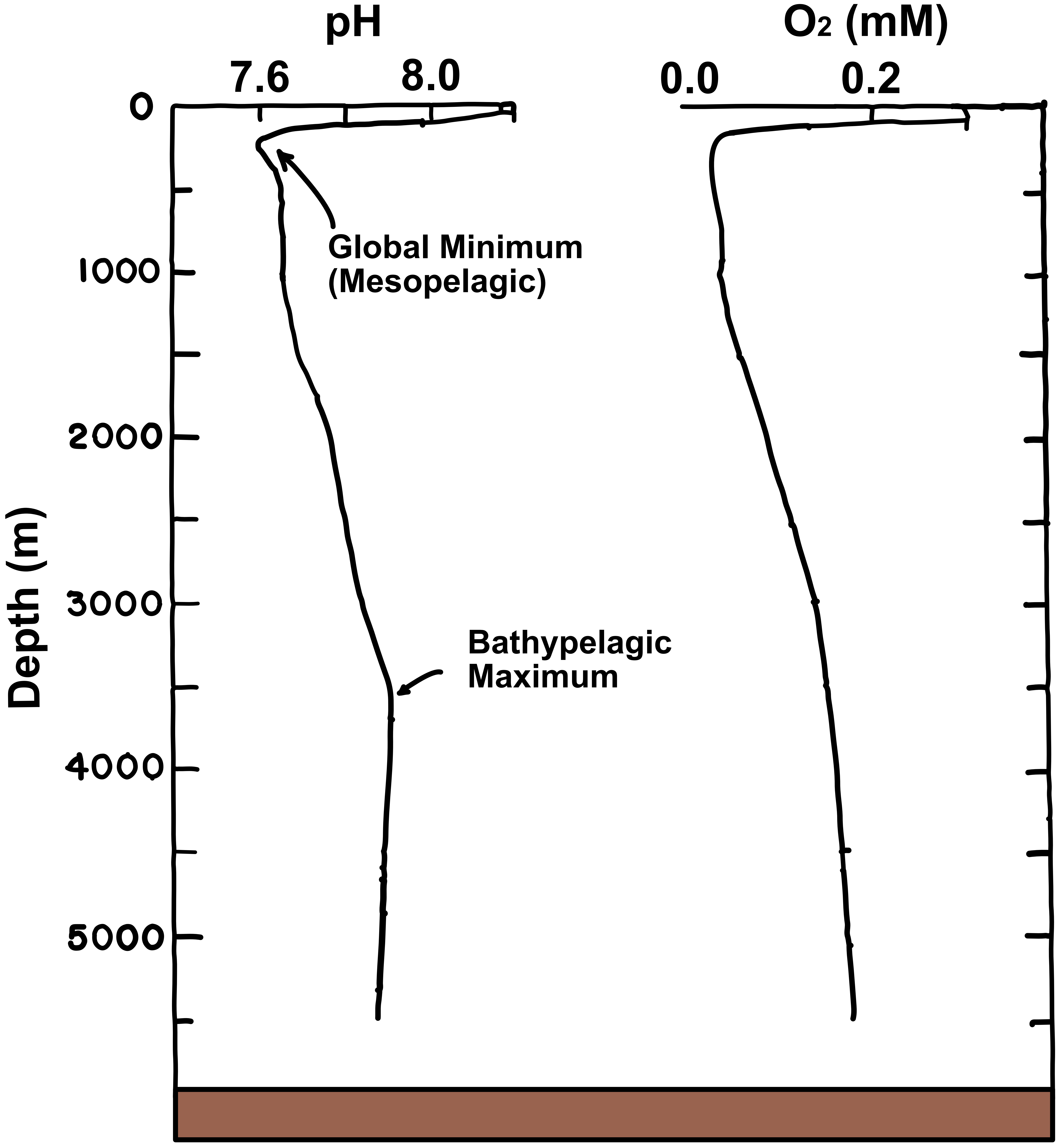

### Figure_S8.pdf

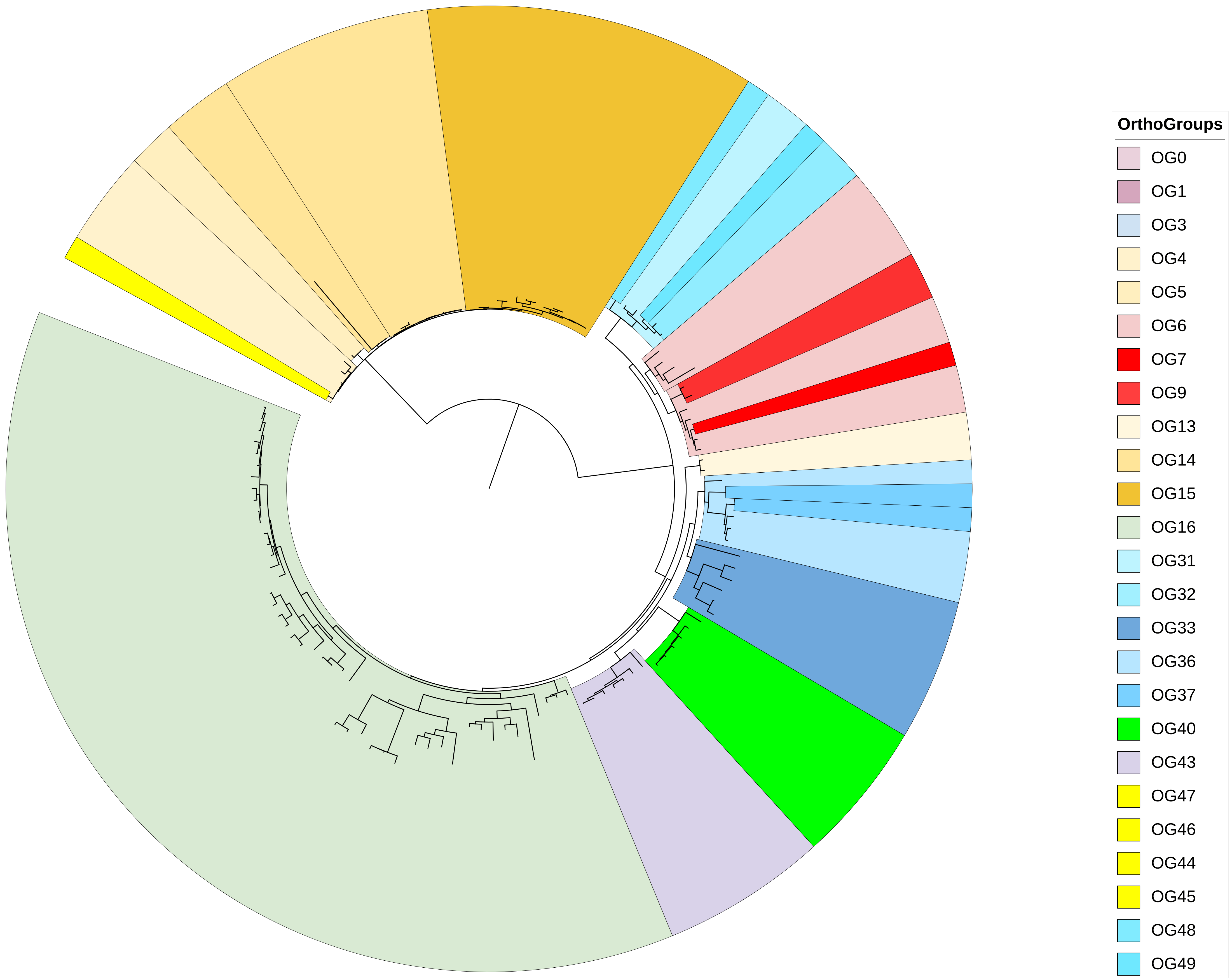

### Figure_S9.1.png

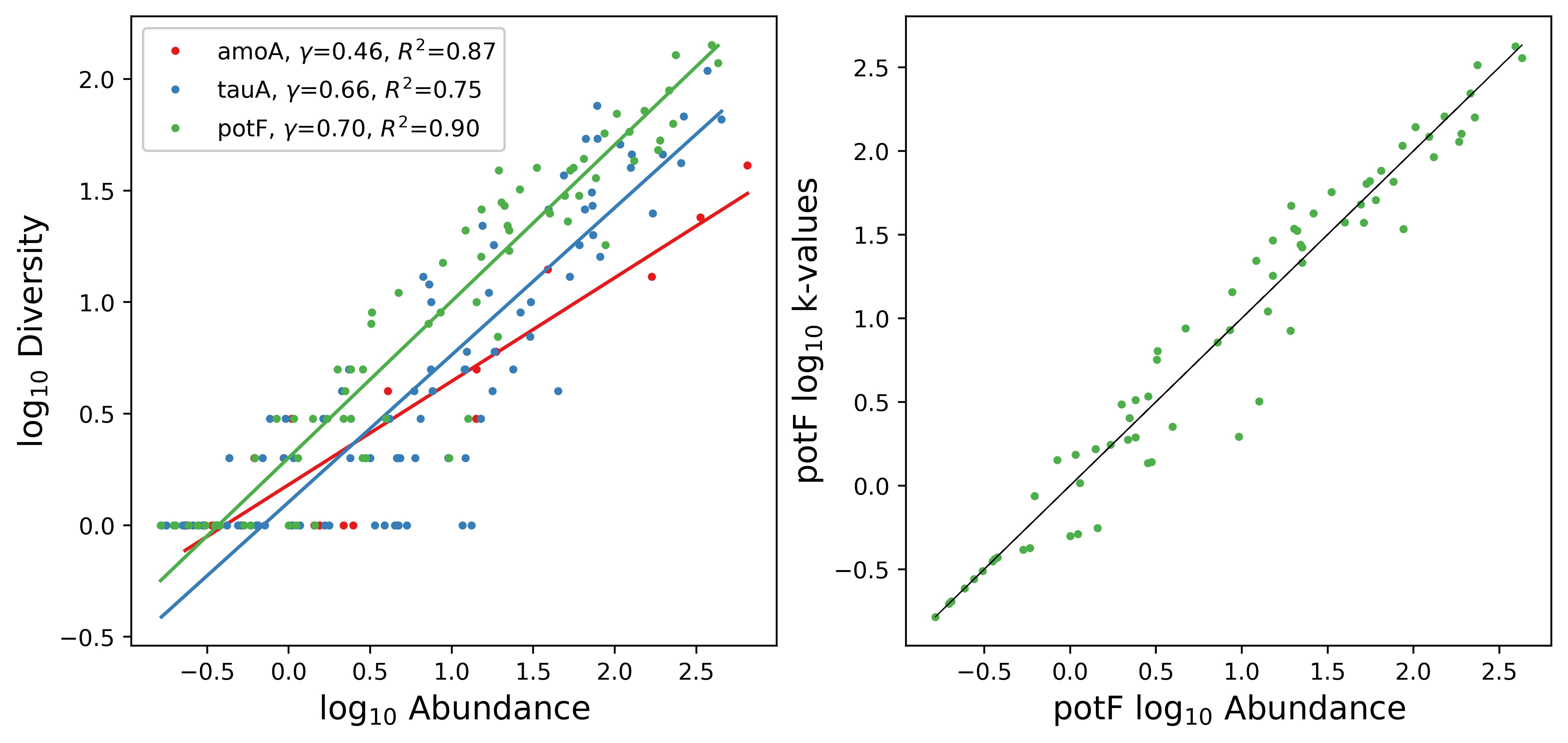

### Figure_S9.2.png

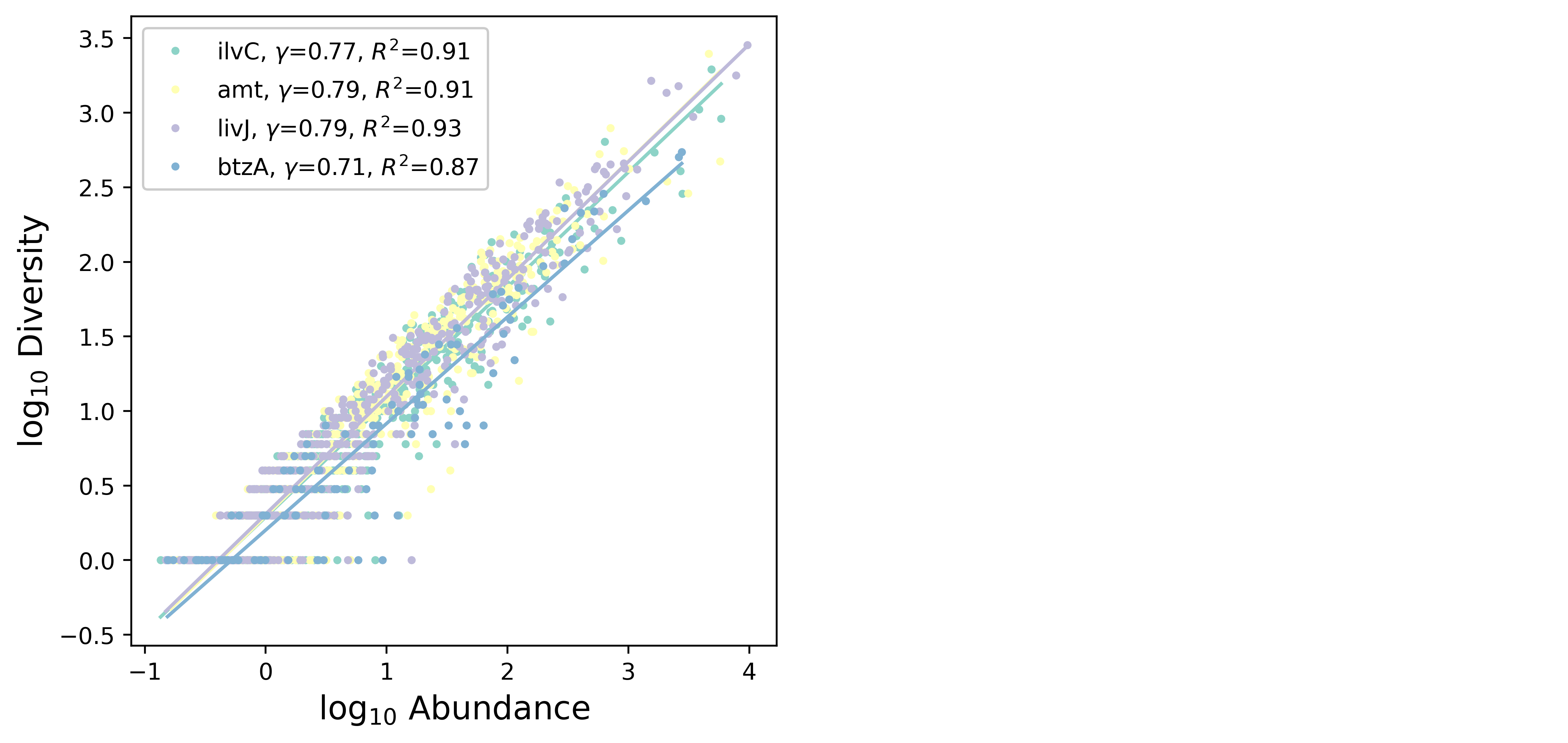
